# A multi-muscular, redundant strategy for free-flight roll stability

**DOI:** 10.1101/2025.09.29.679272

**Authors:** B. Kemper Ludlow, Serene Dhawan, Samuel C. Whitehead, Han Kheng Teoh, Erica Ehrhardt, Noah J. Cowan, Bradley Dickerson, Itai Cohen

**Author notes:** Biological Engineering, California Institute of Technology, USA. Coreva Scientific GmbH & Co KG, Königswinter, Germany.

## Abstract

Whether recovering after a gust of wind, or rapidly saccading away from an oncoming predator, fruit flies show remarkable aerial dexterity about their body roll axis. Here, we investigated the detailed wing kinematic changes during free-flight roll motion and probed the neuromuscular basis for such changes. Consistent with previous work, we observed that flies manipulated the stroke amplitude difference between their wings to control their roll angle. Here, we show that flies are capable of achieving such changes by altering the stroke amplitude of either or both of their wings. Further we found that during corrections flies can also take advantage of an aerodynamically significant change in the angle of attack of their uppermost wing. Curiously, these corrective wing changes cannot be eliminated when motor neurons hypothesized to be used during roll maneuvers (i1, i2, b1, b2, and b3) are individually inhibited. However, free-flight optogenetic manipulations and quasi-steady aerodynamic calculations show that each of these motor neurons individually can effect kinematic changes consistent with a roll correction. Combining this evidence with an analysis of haltere inputs found in the BANC connectome, we propose that the observed robustness could be the result of two sets of muscular redundancies that receive shared inputs from haltere sensory afferents: one set, containing b1 and b2, is able to increase the stroke amplitude of the lower wing; while the other set, containing i1, i2, and b3, is able to decrease the stroke amplitude and wing pitch angle of the upper wing. Because of the redundancy in the input sensory information and output wing motion in the muscles in each cluster, the fly is able to perform roll stability maneuvers even when one of the constituent motor neurons is inhibited. This framework proposes new ways fast aerial maneuverability can be implemented when dealing with the fly’s most unstable rotational degree of freedom.

## Introduction

Flapping flight is a delicate balancing act, requiring organisms to perform constant adjustments to maintain their aerial stability. For the fruit fly, no degree of freedom is as unstable as its body roll angle. Previous analyses have shown flies must begin to correct within 1-2 wingbeats of an external roll perturbation, or risk veering into instability (***Ristroph et al., 2013; Chang and Wang, 2014; Perl et al., 2023***). Remarkably, the fruit fly does just that: whether rolled by a gust of wind in the wild, or an induced perturbation in the laboratory (***Beatus et al., 2015***), its wing motion begins adjusting within just five milliseconds—a single wingbeat—to counteract the perturbation, making this one of the fastest measured stability reflexes (***Beatus et al., 2015***). The speed of this response also makes this reflex an attractive candidate for studying minimal circuits for motor control. Acting on a much faster timescale than the optomotor response time of approximately 50 ms (***Götz et al., 1979; Duistermars et al., 2007a***,b), the roll reflex has been hypothesized to be constrained to a circuit contained entirely within the ventral nerve cord (VNC) (***Figure 1***A). The motor outputs are driven by sensory inputs from the halteres, gyroscopic organs which beat antiphase with the wings and provide rotational velocity as well as timing information to the nervous system via an array of strain sensors spread across their surface (***Pringle, 1948; Dickinson, 1999; Dickerson et al., 2019***). The fast response timing and relative sparseness of the fly nervous system provide upper limits on the total number of neurons which can be used to perform the necessary sensorimotor computations; understanding how this is accomplished could provide insights applicable to motor circuits in other animals and inspire applications in robotics. In addition to being important for stability reflexes, this circuit could also be important for voluntary maneuvers via the control loop hypothesis (***Dickerson et al., 2019***). This hypothesis posits that flies can co-opt their own stability reflexes to steer by manipulating the motion of their own halteres (***Dickerson et al., 2019***). Recent results demonstrating active control of haltere muscles during saccades further support this possibility and make it all the more important to understand the neuromuscular substrate underlying haltere-mediated flight reflexes (***Verbe et al., 2024***).

**Figure 1.**
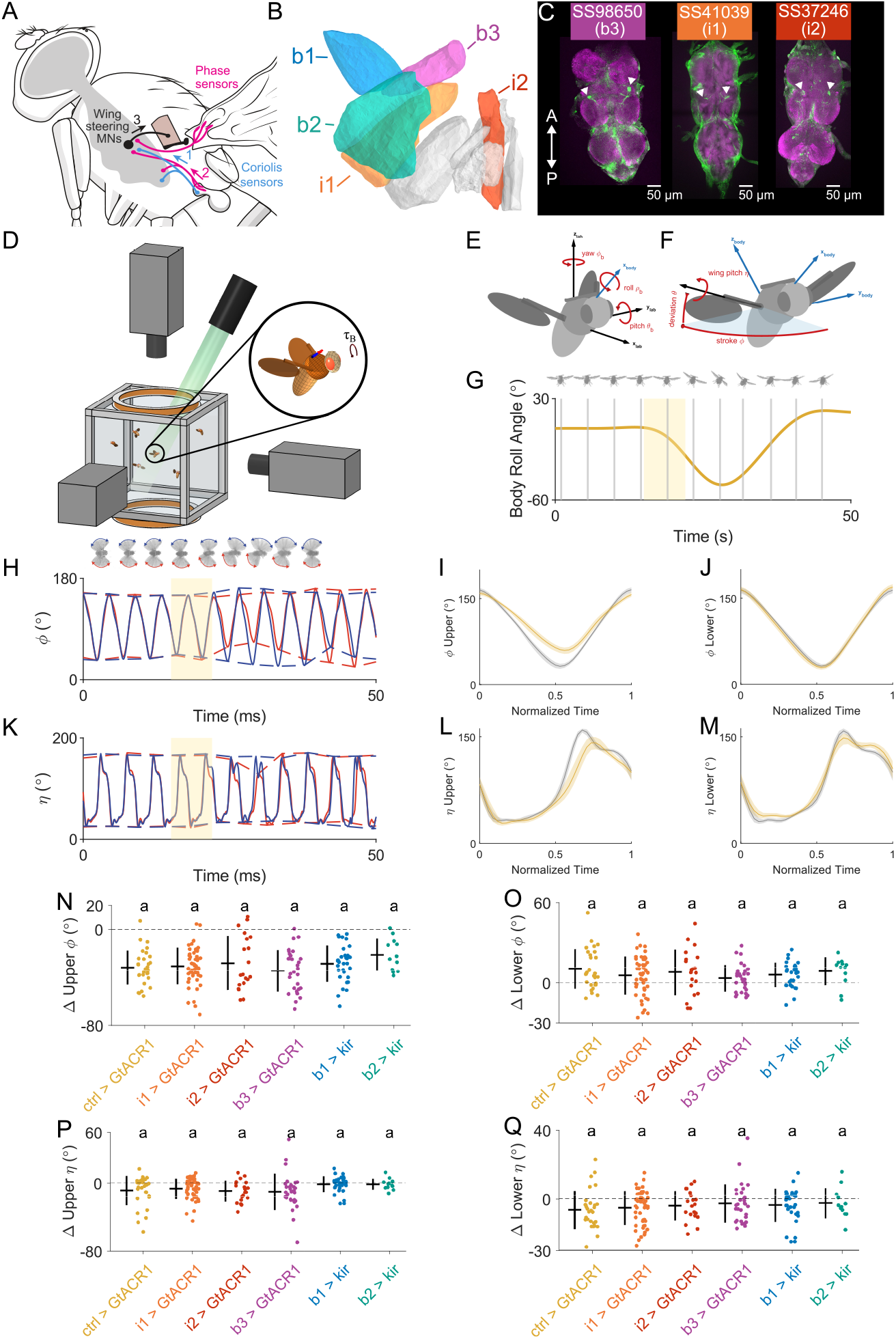
Quantification of roll correction strategies across genotypes. **A** Schematic of halteremediated fast flight control. Sensory information from the wing and halteres is sent to the wing steering muscles to drive changes in wing kinematics. **B** Muscle schematic highlighting five muscles implicated in roll maneuvers: b1 (blue), b2 (teal), b3 (magenta), i1 (orange), and i2 (red). Data are from ***Lindsay et al. (2017)***. **C** Dissections showing expression of GFP driven by split-GAL4 driver lines used to target b3, i1, and i2. **D** Apparatus for inducing roll perturbations. Helmholz coils deliver a magnetic torque to a fly affixed with a magnetic pin, and the response is captured by three highspeed cameras. **E** Definition of body angular degrees of freedom, in the lab frame of reference. **F** Definition of wing angular degrees of freedom, in the body frame of reference. **G** Top: Video stills showing one view of a fly during a roll perturbation. Bottom: Measured body roll angle over time during a roll perturbation. Yellow bar indicates timing of the magnetic impulse. **H** Top: Video stills of the same fly in D, now showing approximate wing stroke amplitude during a roll perturbation. Bottom: Stroke angle of left (blue) and right (red) wings over time during a roll perturbation. **I** Stroke angle of the uppermost wing for a single wingbeat before (grey) and during (yellow) a roll correction, averaged across all movies of genetic control flies (SS01062 > UAS-bGtACR1; N = 17). Shaded regions represent boostrapped confidence interval. **J** As in I, for the lower wing. **K** Wing pitch angle of left (blue) and right (red) wings over time for the roll perturbation shown in D,E. **L** Wing pitch angle of the uppermost wing for a single wingbeat before (grey) and during (yellow) a roll correction, averaged across all movies of genetic control flies. Shaded regions represent boostrapped confidence interval. **M** As in L, for the lower wing. **N** Change in upper wing stroke amplitude during peak correction across all movies with individual motor neurons inhibited (N = 30, 50, 20, 30, 31, and 12, respectively). Data for b1, b2 inhibition are from ***Whitehead et al. (2022)***. **O** As in N, showing change in lower wing stroke amplitude during peak correction. **P** As in N, showing change in peak wing pitch value for the upper wing during peak correction. **Q** As in P, showing change in peak wing pitch value for the lower wing during peak correction. All statistics are Kruskal-Wallis test with Bonferroni correction for multiple hypotheses. **Figure 1—figure supplement 1**. Deviation angle changes during roll perturbations **Figure 1—figure supplement 2**. Controller model fits during roll perturbations

These rapid kinematic changes are implemented by the wing steering system. Unlike the indirect power muscles of the thorax, which provide the main source of lift via thoracic deformations, the steering muscles directly attach to the sclerites of the wing hinge (***Dickinson and Tu, 1997; Balint and Dickinson, 2001***). The activity of these tiny muscles alters the conformation of the wing hinge, thus modulating wing motion and aerodynamic force production (***Heide and Götz, 1996; Tu and Dickinson, 1996; Lehmann and Dickinson, 1998; Balint and Dickinson, 2001, 2004; O’Sullivan et al., 2018; Melis et al., 2024; Ehrhardt et al., 2025***). The steering system includes 12 pairs of direct steering muscles that affix to sclerites in the wing hinge (***Figure 1***B), as well as 5 pairs of indirect steering muscles affixed to other parts of the fly’s thorax. Each muscle is innervated by a single motor neuron and contracts synchronously with the corresponding motor neuron activity (***Balint and Dickinson, 2001, 2004***). Additionally, they are functionally divided: the “tonic” muscles are generally active during flight and contract approximately once per wingstroke, whereas the “phasic” muscles alternate phases of quiescence with bursts of activity (***Heide and Götz, 1996; Lindsay et al., 2017***). Previous work showed that individually silencing the b1 and b2 motor neurons impacted the performance of different parts of the control stabilization model for body pitch (***Whitehead et al., 2022***). The success of this approach raised the question if an analogous set of experiments could be used to probe the faster, even more dynamically unstable roll reflex. Combined, the frameworks for neuromuscular control of pitch and roll would provide a complete picture of how to balance unstable rotational degrees of freedom within a sparse motor system. Here, we performed experiments to investigate the function of the i1, i2, and b3 steering muscles, previously shown to respond to visual roll perturbations (***Lindsay et al., 2017***), through optogenetic manipulation of their motor neurons (***Figure 1***C). Further, we performed a new analysis on data from ***Whitehead et al. (2022)*** to probe more deeply into the effect b1 and b2 have on roll control. We find that the muscles responsible for roll corrections have a control architecture that is distinct from and more robust than that for pitch, indicating the importance of this degree of freedom for flight.

## Results

### Detailed wing kinematics during free-flight roll perturbations

Flies correct for roll perturbations primarily by inducing a difference between their left and right stroke amplitudes (***Dickinson et al., 1999; Beatus et al., 2015***). Activity in i1, i2, and b3 is correlated with a decrease in stroke amplitude, which would be one possible way to induce this difference (***Ehrhardt et al., 2025; Melis et al., 2024***). At the same time, activity in b1 and b2 has been correlated with increasing stroke amplitude, which could also generate the necessary asymmetry (***Whitehead et al., 2022; Ehrhardt et al., 2025; Melis et al., 2024***). These studies raise several questions including what is the relative importance of increasing or decreasing stroke amplitudes during recovery from roll perturbations, and which muscles are responsible for these changes. To answer these questions, we began with a detailed analysis of wing kinematic during roll perturbations on genetic controls, and then applied these same analyses to experiments with inhibition of one of our candidate motor neurons (***Figure 1***C).

We recorded the behavior of freely-flying *Drosophila* while applying an external magnetic field pulse which acts on small magnetic pins glued onto the backs of the flies to induce roll perturbations (***Figure 1***D). Simultaneously, we co-triggered a collimated LED to apply optogenetic silencing or activation during the same time period. From these experiments, we extracted the full wing and body kinematics of the flies’ motion (as defined in ***Figure 1***E,F). An example trajectory of body roll over time during one of these perturbations is shown in ***Figure 1***G. Here, the optogenetic LED was on from t = 0 to 50 ms, and the fly experienced a magnetically driven roll torque during the time period highlighted in yellow and recovered its roll orientation within ∼5 wingbeats. We present the corresponding extracted wing stroke angle versus time for the left (blue) and right (red) wings in ***Figure 1***H. When the fly corrected for the roll perturbation, it did so by inducing an asymmetry in the stroke amplitude of its left and right wings, as previously reported by ***Beatus et al. (2015)***. Specifically, it did so by decreasing the stroke amplitude of the wing that is in the uppermost position immediately following the perturbation (or contralateral to the direction of the roll perturbation— in this case, the right wing, in red), and simultaneously increasing the stroke amplitude of the lower wing (or ipsilateral to the direction of the roll perturbation—left wing, blue). On average, across all roll perturbations of genetic controls, we found a large, significant decrease in the upper stroke amplitude, as well as a smaller but still significant increase in the lower stroke amplitude (***Figure 1***I-J). The extracted wing pitch angles for the left (blue) and right (red) wings are shown in ***Figure 1***K. Here, we also observed a decrease in the peak wing pitch angle on the upper wing, which corresponds to a steeper angle of attack when the wing is sweeping from back to front. Such changes in the angle of attack can induce aerodynamic torques that may also affect roll. This decrease in the peak wing pitch angle is generally conserved across roll perturbations (***Figure 1***L). We also observed a small overall decrease in the amplitude of the wing pitch angle on the lower wing (***Figure 1***M). Collectively, we observed several changes to the wing stroke amplitude and angle of attack with the potential to effect a roll response (See ***Figure 1***—***figure Supplement 1*** for changes to the deviation angle, which we found are not aerodynamically significant).

To investigate the underlying neuromuscular mechanism for enacting these wing kinematic changes and ultimately correcting for roll, we sought to quantify the individual effects of the i1, i2, b1, b2, and b3 muscles, shown in ***Figure 1***B, on each of these four correction strategies: decreasing upper stroke amplitude, increasing lower stroke amplitude, decreasing upper wing pitch peak angle, and decreasing lower wing pitch amplitude. We used previously developed split-GAL4 driver lines (***Ehrhardt et al., 2025; Cheong et al., 2024***) that target i1, i2, and b3 motor neurons separately to perform free-flight roll perturbation experiments while optogenetically silencing each of these motor neurons (***Figure 1***C). Additionally, we performed further analysis on data from ***Whitehead et al. (2022)*** in which roll perturbations were applied while the b1 or b2 motor neurons were chronically inhibited using Kir2.1. We found that no matter which motor neuron was inhibited, flies still decreased their upper stroke amplitude by similar amounts in response to roll perturbations when compared with controls (***Figure 1***N). Likewise, the distributions of lower stroke amplitude changes were similar across all conditions (***Figure 1O***). For wing pitch, we also observed comparable distributions of change in peak wing pitch angle for both wings across conditions, with a wide range of values consistent with our genetic controls (***Figure 1***P,Q).

In addition to examining detailed wing kinematic changes during roll corrections, we also applied a proportional-integral (PI) model for roll stability developed in ***Beatus et al. (2015)***. In this model, the fly’s roll velocity 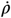 is taken as an input signal which is then transformed to give a prediction of wing kinematic output—in this case, the difference between left and right stroke amplitude ΔΦ_*LR*_—using the equation

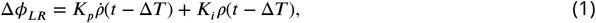

where the proportional gain *K*_*p*_, integral gain *K*_*i*_, and time delay Δ*T* are fit to the experimental data. We found that across silencing conditions, there was no significant change in any of the model fit parameters (***Figure 1***—***figure Supplement 2***). Our kinematic and controller analyses thus indicate that individually inhibiting any of the motor neurons corresponding to muscles implicated in roll control does not impede the fly’s ability to maintain its body roll angle. These results are somewhat surprising given the overall sparseness of the fly steering system and prior results that show they are active during roll maneuvers (***Lindsay et al., 2017***).

### Free-flight optogenetic manipulation of i1, i2, and b3

In search of clues about how the motor system could maintain its robust roll control without access to seemingly crucial steering muscles, we investigated how these muscles affect flight when manipulated during free flight in the absence of a mechanical perturbation. Previous results from free flight experiments have shown that excitation of the b1 or b2 motor neurons increases the wing stroke amplitude (***Whitehead et al., 2022***). In addition, results from tethered experiments suggest that excitation of the i1, i2, or b3 motor neurons decreases the stroke amplitude (***Ehrhardt et al., 2025; Melis et al., 2024***). To reveal the effect of i1, i2, or b3 motor neuron excitation in free flight, we optogenetically stimulated flies expressing UAS-CsChrimson or UAS-GtACR1 driven by our selected driver lines to activate or inhibit the selected neurons in the absence of a mechanical perturbation, and recorded changes in body angular motion and wing kinematics. ***Figure 2*** shows the resulting body and wing motion from optogenetic activation of the i1 and i2 motor neurons, as well as optogenetic inhibition of the b3 motor neuron (See ***Figure 2—figure Supplement 1*** showing results from inhibition of i1 and i2, as well as activation of b3, which showed no significant effect on the flies’ wing or body motions). When activating the i1 motor neuron, we observed flies pitch steeply down, while body roll and yaw angles remained mostly unchanged (***Figure 2***A). This change seems to be primarily driven by a large decrease in stroke amplitude (***Figure 2***B); additionally we observed a decrease in the peak and amplitude of the wing pitch angle (***Figure 2***C). In the most extreme responses to activation, the stroke amplitude decreased to zero, and the flys’ wings stalled in mid-air (See ***Figure 2***—***video 1***). Activating the i2 motor neuron yielded very similar results: a drop in body pitch angle likely caused by decreasing stroke amplitude, as well as a decrease in the peak and amplitude of the wing pitch angle (***Figure 2***D-F). Inhibiting b3 produced results in the opposite direction; flies pitched slightly up while increasing their stroke amplitudes (***Figure 2***G,H).

**Figure 2.**
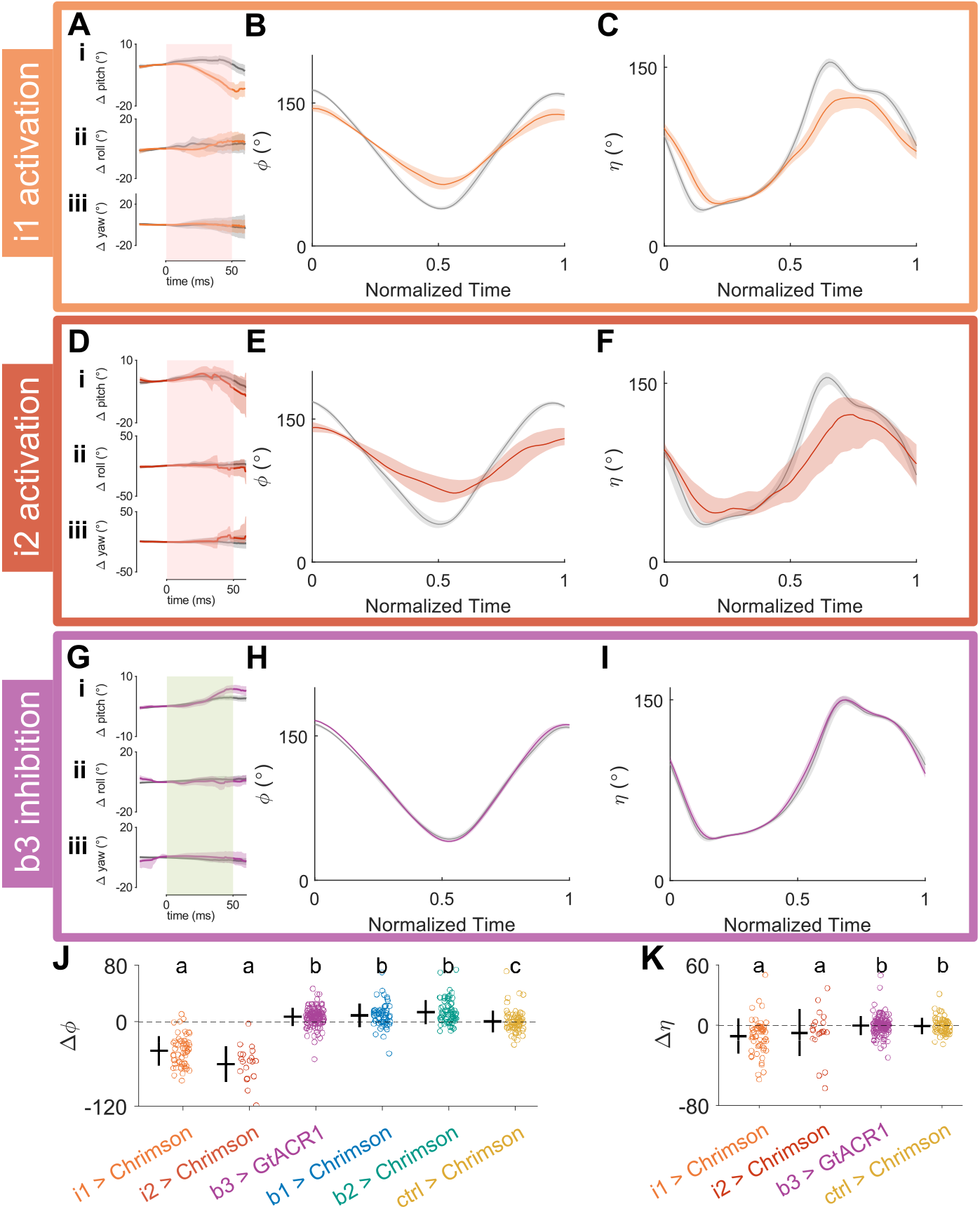
Free-flight optogenetic manipulation. **A** Average body pitch (i), roll (ii), and yaw (iii) during optogenetic activation of the i1 motor neuron (SS41039 > UAS-CsChrimson, orange, N = 119) compared to genetic controls (SS01062 > UAS-CsChrimson, grey, N = 195). Red bar indicates timing of optogenetic stimulus. Shaded areas represent bootstrapped confidence interval. **B** Stroke angle of for a single wingbeat before (grey) and during (orange) optogenetic activation of the i1 motor neuron, averaged across all movies. **C** Wing pitch angle for a single wingbeat before (grey) during (orange) optogenetic activation of the i1 motor neuron, averaged across all movies. **D-F** As in A-C, for optogenetic activation of the i2 motor neuron (SS37246 > UAS-CsChrimson, red, N = 38). **G-I** As in A-C, for optogenetic inhibition of the b3 motor neuron (SS98650 > UAS-GtACR1, purple, N = 139. Genetic controls SS01062 > UAS-GtACR1, grey (A i-iii), N = 557). **J** Change in stroke amplitude across all free-flight opotogenetic experiments. (N = 195, 119, 38, 139, 75, and 84 respectively). Data for b1, b2 activation are from ***Whitehead et al***. (***2022***). **K** As in J, for the change in peak wing pitch angle. All statistics are Kruskal-Wallis test with Bonferroni correction for multiple hypotheses. **Figure 2—figure supplement 1**. Optogenetic inhibition of i1, i2 and activation of b3 **Figure 2—video 1**. Example video showing wings stalling during i1 activation using CsChrimson.

A comparatively small effect on the wing pitch angle was observed from b3 inhibition (***Figure 2***I). These kinematic changes are summarized in ***Figure 2***J,K, along with the changes in stroke amplitude during b1 and b2 activation first reported in ***Whitehead et al. (2022)***.

Whereas the results from direct activation of i1 and i2 are relatively straightforward to interpret, the results from b3 inhibition require a bit more context. Anatomically, b3 is attached to the same sclerite as b1 and b2, which can increase the stroke amplitude, but pulls antagonistically against them (***Dickinson and Tu, 1997***). Thus, when b3 is inhibited, the stroke amplitude is most likely increased by preexisting activity in b1 and b2 (***Whitehead et al., 2022; Melis et al., 2024***). Interestingly, activating b3 in free flight did not significantly shorten the wing stroke amplitude (***Figure 2***—***figure Supplement 1***). We hypothesize that it may have been overpowered by existing activity in b1 and/or b2. In this picture, during maneuvers when activity in b1 or b2 is suppressed, b3 activity would cause a shortening of the wingstrokes. Indeed, this pattern of coordinated activity has been observed in tethered flight (***Lindsay et al., 2017***). This picture is also consistent with the observation that b3 activity is correlated with a decrease in wingstroke amplitude (***Melis et al., 2024***). These results indirectly demonstrate that, much like i1 and i2, the b3 muscle can act to shorten the wingstroke.

Collectively, our free-flight optogenetic experiments show that these muscles appear to be clustered in their function for changing the wing stroke: i1, i2, and b3 all act to reduce the stroke amplitude or prevent it from increasing further, while b1 and b2 act to increase the stroke amplitude. Additionally, i1 and i2 seem to affect the wing pitch angle in ways consistent with changes seen during roll perturbations.

### Quasi-steady aerodynamic analysis of roll torque generation

The wing kinematic changes observed during optogenetic manipulation of the i1, i2, b3, b1, and b2 motor neurons appear qualitatively similar to those changes observed during a roll correction. To confirm that they could also generate the necessary aerodynamic torques, we performed a series of quasi-steady aerodynamic aerodynamic analyses. First, we analyzed the kinematic changes when genetic control flies correct for a roll perturbation, as in ***Figure 1***D-L. To do so, we selected a candidate experiment in which a fly was observed to perform a roll correction without performing any extraneous maneuvers before or after the perturbation (***Figure 3***A). We isolated the kinematics of one wingbeat before the perturbation (***Figure 3***A, grey box) and the wingbeat of peak corrective activity (***Figure 3***A, yellow box). We then linearly interpolated the kinematic profiles for each rotational degree of freedom on each wing (three degrees of freedom × two wings, for six variables total) between the pre-perturbation and maximally corrective wingbeats, generating 50 kinematic profiles for each variable (***Figure 3***B,i-vi). This allowed us to select profiles for each variable to “mix and match” how much each corrective strategy is used. We then used quasi-steady aerodynamic calculations, based on prior work in ***Whitehead et al. (2022, 2015); Wang et al. (2004); Sane and Dickinson (2002); Dickinson et al. (1999)***, to compute the average torque over one wingbeat using a particular set of kinematic profiles. We found it most useful to isolate one corrective strategy and vary the corresponding kinematic profile while holding the other five profiles at a constant percentage between pre-perturbation and corrective values. Using this technique, we first calculated aerodynamic torques about the fly’s body roll axis, as diagrammed in ***Figure 3***C.

**Figure 3.**
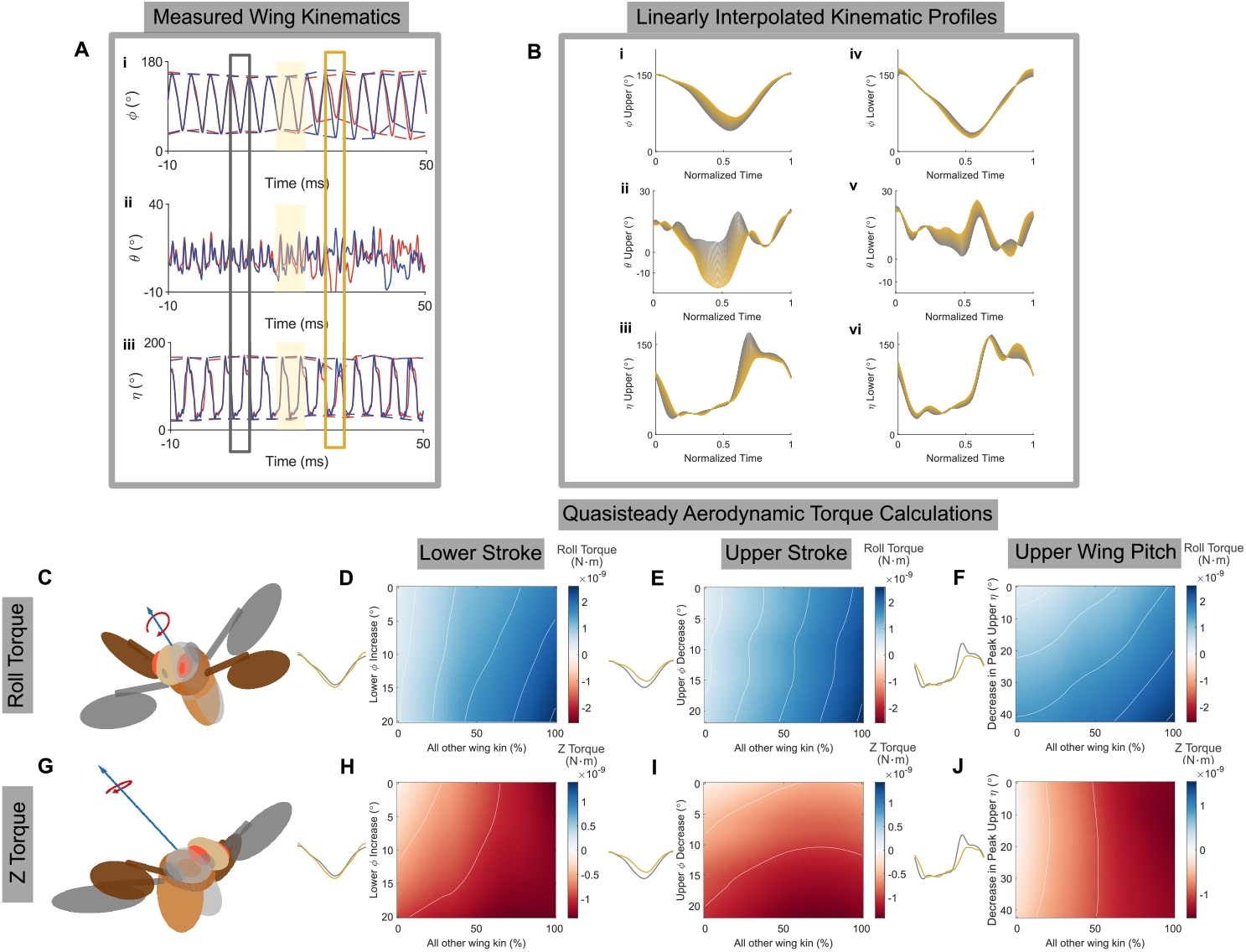
Quasi-steady aerodynamic calculations to determine relevance of kinematic changes to corrective torque. **A** Experimentally measured stroke (i), deviation (ii), and wing pitch (iii) angles during a roll correction. **B** 50 Linearly interpolated wing kinematic profiles for a single wingbeat, ranging from before perturbation (grey) to the maximally corrective wingbeat (yellow): (i) upper wing stroke angle, (ii) upper wing deviation angle, (iii) upper wing pitch angle, (iv) lower wing stroke angle, (v) lower wing deviation angle, and (vi) lower wing pitch angle. **C** Schematic of rotation about body roll axis correcting a perturbed fly (grey) to zero body roll angle (color). **D** Wingbeat-averaged roll torque generated by different combinations of the wing stroke profiles in B. Y axis varies only lower wing stroke amplitude from pre-perturbation (grey) to maximally corrective (yellow). X axis uniformly varies all other wing kinematic profiles from pre-perturbation (grey) to maximally corrective (yellow). **E** As in D, with upper stroke profile varying along the y axis and all other kinematic profiles varying along the x axis. **F** As in D, with upper wing pitch profile varying along the y axis and all other kinematic profiles varying along the x axis. **G** Schematic of rotation about body z axis correcting a perturbed fly (grey) to zero body roll angle (color), at the cost of slightly altered pitch and yaw angles. **H-J** As in E-G, with wingbeat-averaged z torque plotted. All contour lines are spaced by 5 × 10^−10^ N⋅m. **Figure 3—figure supplement 1**. Visualization of changing only upper wing pitch during wingstroke **Figure 3—figure supplement 2**. Quasi-steady aerodynamic calculations for other wing kinematic variables

In ***Figure 3***D, the amount of lower wingstroke amplitude increase is varied along the y axis, while the amount of change in the other five kinematic variables is varied uniformly along the x axis. Each point on the heatmap represents the average body roll torque for one combination of kinematic profiles. As the lower stroke amplitude increased from pre-perturbation to peak correction levels, we found little impact on the roll torque compared with the changes from the other five kinematic variables. Similarly, decreasing the upper stroke amplitude did not significantly increase the roll torque compared with the changes to the other five kinematic variables (***Figure 3***E). However, decreasing the upper wing pitch angle *did* contribute more significantly to the overall roll torque during the correction, shown by the increasing torque value when moving down the y axis in ***Figure 3***F. The effect of this change in wing pitch is diagrammed in ***Figure 3—figure Supplement 1***; the angle of attack remains more vertical after the dorsal wing flip, and so the force perpendicular to the wing points less upward, and less roll torque is generated on that side of the body. Results for the other three kinematic variables, which were not found to produce significant corrective torques, are shown in ***Figure 3***—***figure Supplement 2***. These calculations lead us to conclude that the key contributor to roll torque in this perturbation was the fly’s decrease in peak upper wing pitch angle.

However, when a fly is perturbed about its body roll axis, there is a secondary axis it can rotate around to recover its previous roll angle. This is due to the fact that rotations about one principal axis can move the other principal axes, and different sequences of rotations can produce similar results depending on the order in which they are performed. As previously reported by ***Beatus et al. (2015)***, counter rotations about the body z axis, which is perpendicular to the roll axis and extends out through the top of the fly’s thorax, can recover body roll angle at the cost of a smaller change in pitch and yaw angles (***Figure 3***G). This mechanism give flies another axis about which they can generate torque to recover their body roll angle. Additionally, the change in yaw heading may be beneficial for correcting heading changes caused by a roll perturbation. Using the same strategy shown in ***Figure 3D-F***, we evaluated the impact of each of our three identified corrective strategies on z torque. This time, we found that increasing lower stroke amplitude (***Figure 3***H) and decreasing upper stroke amplitude (***Figure 3***I) both increased the amount of z torque. Conversely, decreasing the peak wing pitch value had little to no effect on z torque generation (***Figure 3***J). From these calculations, we conclude that the observed wing kinematic strategies allow flies to efficiently combine roll and z torque when recovering from mid-flight stumbles by leveraging multiple changes over their wingstroke.

Finally, we applied a similar analysis to our optogenetic data to evaluate the effectiveness of wing kinematic changes driven by our muscles of interest in generating corrective roll and z torques. To do so, we started by selecting an example movie from each optogenetic condition and isolated one wingbeat before and during optogenetic stimulation. An example for activation of the i1 motor neuron is shown in ***Figure 4***A, with the pre-stimulus wingbeat highlighted in grey, and stimulus wingbeat highlighted in orange. We then linearly interpolated between pre-stimulus and stimulus kinematics on only one wing (***Figure 4***B,i-iii), while keeping the other wing’s kinematics fixed at pre-stimulus trajectories (***Figure 4***B,iv-vi). Doing so allowed us to gradually introduce stimulation kinematics on only one side of the body and generate asymmetries similar to those observed during roll corrections. Because our optogenetic manipulations are symmetric, we could have designated either wing as the upper or lower wing. In ***Figure 3*** we used an example video in which the right wing was the upper wing. For consistency, when we analyzed an optogenetic manipulation movie in which the wing pitch or stroke amplitude decreased, we applied these changes to the right wing while holding the left wing kinematics fixed (***Figure 4***C-J). Conversely, when we analyzed a movie in which the stroke amplitude increased, we applied these changes to the left wing while holding the right wing kinematics fixed (***Figure 4***K-P). This procedure ensured that the directions of the modeled roll and z torques stayed consistent across the analyses. This approach differs slightly from that in ***Figure 3***, as we now compared non-perturbed to optogenetically perturbed wing kinematics, rather than non-corrective to corrective wing kinematics. Nevertheless, this analysis provides insight into how the kinematic changes driven by various muscle activations can contribute to corrective behavior.

**Figure 4.**
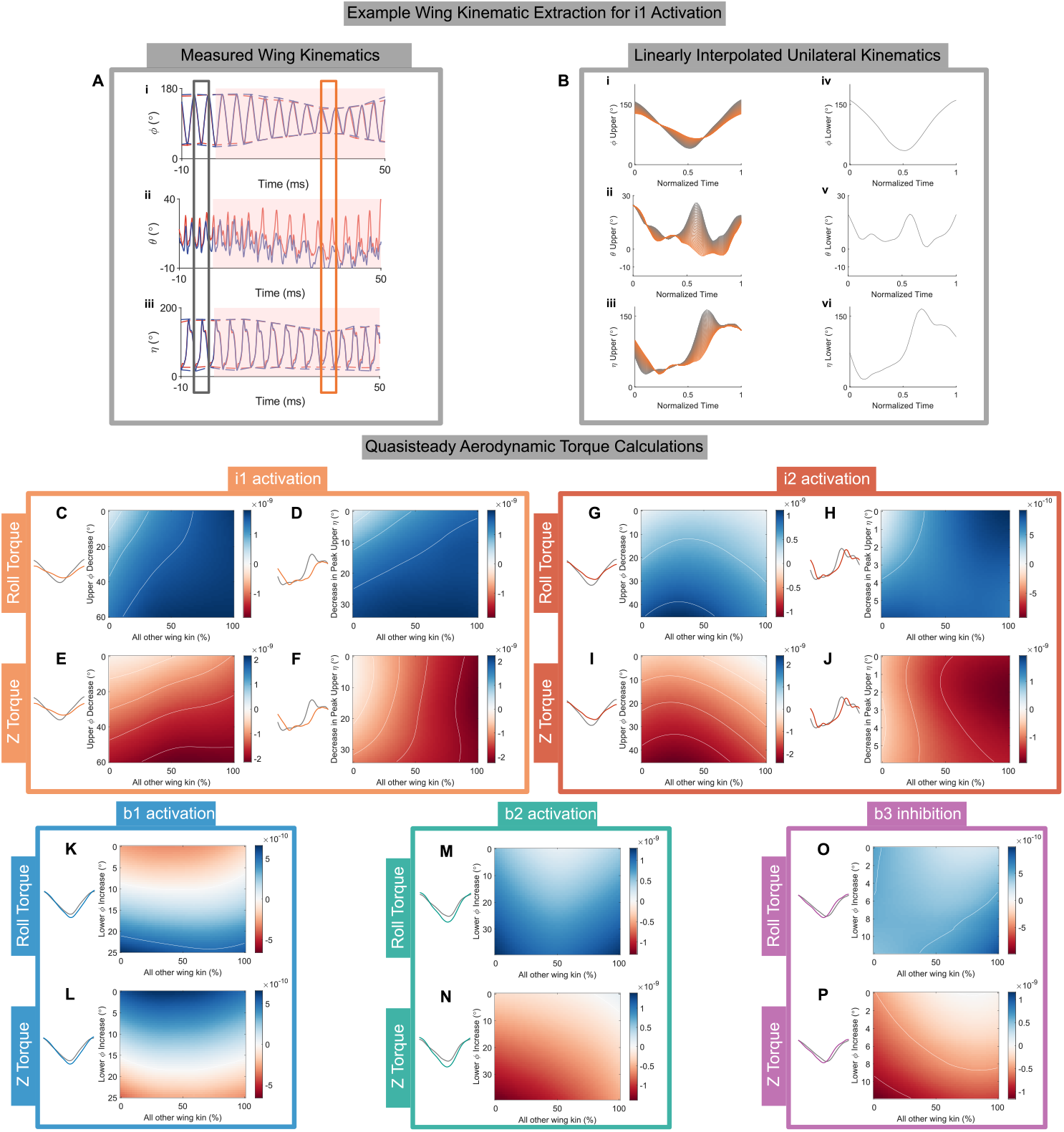
Quasi-steady aerodynamic calculations simulating the effect of wing kinematic changes caused by optogenetic manipulation on aerodynamic torque when applied unilaterally. **A** Experimentally measured stroke (i), deviation (ii), and wing pitch (iii) angles during optogenetic activation of the i1 motor neuron. **B** 50 Linearly interpolated wing kinematic profiles for a single wingbeat, ranging from before stimulus (grey) to approximately 30 ms after i1 activation (orange) on one wing, while the other wing kinematic profiles are kept static (pre-stimulus kinematics, grey): (i) “upper” (right) wing stroke angle, (ii) “upper” (right) wing deviation angle, (iii) “upper” (right) wing pitch angle, (iv) “lower” (left) wing stroke angle, (v) “lower” (left) wing deviation angle, and (vi) “lower” (left) wing pitch angle. **C** Wingbeat-averaged roll torque generated by different combinations of the wing stroke profiles in B. Y axis varies only “upper” (right) wing stroke amplitude from pre-stimulus (grey) to optogenetically activated (orange). X axis uniformly varies other “upper” (right) wing kinematic profiles from pre-stimulus (grey) to optogenetically activated (orange). “Lower” (left) wing kinematics never change. **D** As in C, with “upper” (right) wing pitch angle varying along the y axis instead. **E**,**F** As in C,D, with wingbeat-averaged z torque plotted instead. **G-J** As in D-G, using kinematic profiles for i2 optogenetic activation. **K** As in C, using kinematic profiles for b1 optogenetic activation with “lower” (left) wing stroke angle varying along the y axis, and other “lower” (left) wing kinematic profiles varying along the x axis. “Upper” (right) wing kinematics never change. **L** As in K, with wingbeat-averaged z torque plotted instead.**M**,**N** As in K,L, using kinematic profiles for b2 optogenetic activation. Raw data for b1, b2 activation are from ***Whitehead et al.(2022***). **O**,**P** As in M,N, using kinematic profiles for b3 optogenetic inhibition. All contour lines are spaced by 5 × 10^−10^ N⋅m. **Figure 4—figure supplement 1**. Linearly interpolated profiles for i2 activation **Figure 4—figure supplement 2**. Linearly interpolated profiles for b1 activation **Figure 4—figure supplement 3**. Linearly interpolated profiles for b2 activation **Figure 4—figure supplement 4**. Linearly interpolated profiles for b3 inhibition

In ***Figure 4C***, the decrease in stroke amplitude on the right wing due to i1 activation is varied along the y axis, while the changes in deviation and wing pitch angle on the right wing due to i1 activation are varied along the x axis. We found that the observed decrease in stroke amplitude can contribute to roll torque, but a larger contribution came from the decrease in peak wing pitch angle (***Figure 4D***). The observed decrease in stroke amplitude did, however, contribute the majority of z torque (***Figure 4E***), while very little z torque was generated by the decrease in peak wing pitch angle (***Figure 4F***). In the case of i2 activation, we found that the stroke amplitude decrease also contributed to the total roll torque (***Figure 4G***), but in this case the observed changes in wing pitch angle, while able to generate some roll torque, contributed to a lesser magnitude than the stroke amplitude changes (***Figure 4H***). Consistent with our results for roll perturbations and i1 activation, the stroke amplitude decrease caused by i2 activation can drive a large increase in z torque (***Figure 4I***), but the corresponding changes in wing pitch angle have little to no effect on the z torque (***Figure 4J***).

Next we turned to the analysis of experiments in which the optogenetic stimulus increased the stroke amplitude, as would be observed in the lower wing during a roll correction. While activation of b1 and b2, and inhibition of b3, were all observed to increase wing stroke amplitude, the magnitude of this increase is smaller than the magnitude of the decrease in stroke amplitude when i1 and i2 are activated. Consequently, the torques these wing stroke increases were predicted to produce were about 2 to 4 times smaller. We found that the stroke amplitude increase caused by b1 activation increased both roll and z torque (***Figure 4K***,L). The stroke amplitude increase caused by b2 activation showed little effect on roll or z torques when implemented in isolation from the other wing kinematics (***Figure 4M***,N). Finally, the stroke amplitude increase during b3 inhibition appeared to have very little effect on roll torque (***Figure 4O***), but did slightly increase z torque (***Figure 4P***).

Our calculations show the individual contributions of optogenetic muscle activations (or inhibition, in the case of b3) can vary greatly. We see many consistencies, however, with the corrective torques observed during roll perturbations: decreasing the upper stroke amplitude drives z torque in the case of i1 and i2 activation, decreasing the peak upper wing pitch angle drives roll torque in the case of i1 activation, and increasing the lower stroke amplitude can increase z torque in the case of b1 activation, as well as b3 silencing. Thus, not only are the torque contributions consistent, but we also note that in the cases of the observed stroke amplitude changes, there are multiple muscles that can enact each change. These results support a hypothesis that muscles with similar kinematic outputs may be redundantly controlled together, which would allow to fly to maintain roll stability while one of the corresponding motor neurons is silenced.

### Connectome evidence for redundancy in roll control

The conclusions from our free-flight results are further supported by an analysis of the fly connectome. Here, we focus on what inputs these motor neurons receive from the haltere sensory system, the mechanosensory organs which provide the fly with information about its own rotational velocity (***Pringle, 1948; Hengstenberg, 1988; Dickinson, 1999***). Prior results on the Female Adult Nerve Cord (FANC) dataset showed that haltere afferents are organized into distinct morphological clusters, which target different subsets of wing steering motor neurons (***Dhawan et al., 2025; Phelps et al., 2021***). We repeated these analyses on the Brain and Nerve Cord (BANC) dataset to further confirm their validity and investigate how these clusters could contribute to robust roll control. Both the FANC and BANC datasets used female flies, so we can compare them without needing to account for any sexual dimorphism in the wing motor system that may arise from to its use in male courtship song (***O’Sullivan et al., 2018***). This approach is also consistent with our free-flight experiments, which only use female flies due to their more robust flight behavior (see Methods and Materials). First, we identified reconstructions of the haltere sensory afferents as well as the b1, b2, i1, i2, and b3 motor neurons within the BANC dataset (***Figure 5A***). We then identified all direct connections from haltere afferents to the steering motor neurons of interest, shown in ***Figure 5B***. We observed clustering of the haltere inputs to particular motor neurons. We quantified this clustering by performing a cosine similarity analysis on the synaptic inputs from the haltere to the wing steering motor neurons (***Figure 5C***). We found that the motor neurons can be grouped into two distinct clusters based on their inputs: one containing b1 and b2, the other containing i1, i2, and b3. Although i2 does not appear to be included in the same cluster as i1 and b3 when looking at the data for the right hand side of the fly, this asymmetry is believed to be due to errors in the machine learning-automated segmentation process, as the left hand side data and previous results from the FANC data set indicate i2 is part of the i1, b3 cluster (***Dhawan et al., 2025; Lesser et al., 2024***). For the specific case of roll control, pathways that carry haltere sensory information across the midline may also be particularly important for driving the asymmetric changes seen during corrective maneuvers. Interestingly, such pathways show similar clustering to the direct haltere sensory inputs. ***Figure 5D*** shows an example pathway that utilizes the wing contralateral haltere interneurons (wCHINs, ***Trimarchi and Murphey (1997); Strausfeld and Seyan (1985)***) to carry sensory information from the left side of the body to the i1, i2, b3 cluster on the right side (red), compared with the direct pathway carrying sensory information from the left hand side haltere afferents to the left hand side steering muscle cluster (blue). The picture that emerges from this analysis is that the b1 and b2 motor neurons are receiving one set of excitatory inputs whereas the i1, i2, and b3 motor neurons are receiving a separate set of inputs. Importantly, since there are two or more motor neurons in each cluster, this architecture could in principle also explain the robust function under inhibition of individual motor neurons.

**Figure 5.**
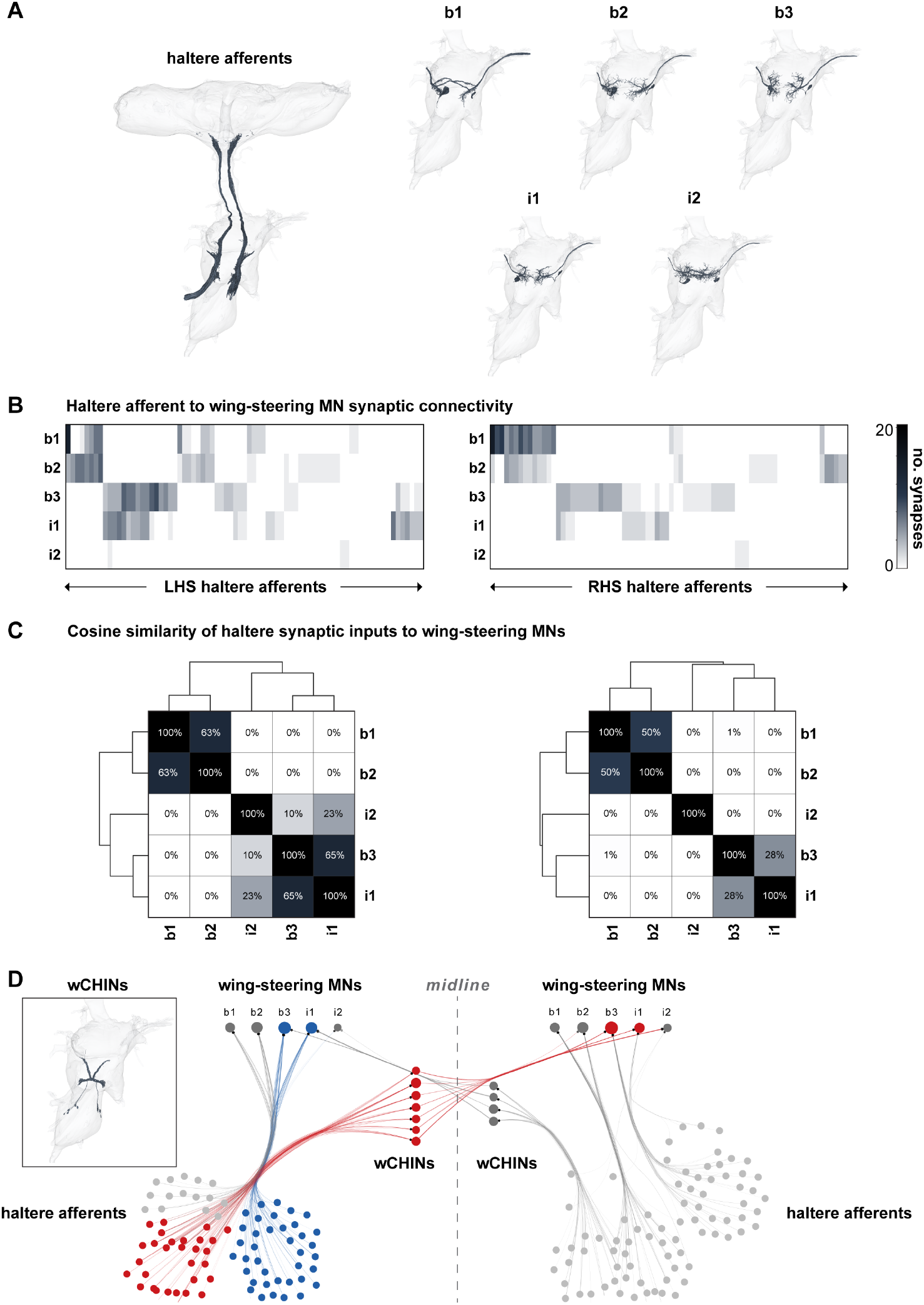
Connectome analysis using BANC dataset. **A** EM renderings of haltere afferents and b1, b2, b3, i1, and i2 motor neurons. **B** Synapses on LHS (left) and RHS (right) of haltere afferents onto selected motor neurons. **C** Cosine similarity of normalized haltere inputs on LHS (left) and RHS (right). **D** Example synaptic pathway (red) through wCHINs (reconstruction in inset) across the midline, compared to direct connectivity cluster (blue)

## Discussion

Collectively, the results of our experiments illustrate that fast, robust roll control is achieved through redundancies within and across two distinct muscle groups. Specifically, our experiments demonstrated that flies use three distinct strategies for roll correction: decreasing the upper wing stroke amplitude; increasing the lower wing stroke amplitude; and modifying the upper wing pitch angle. These changes were conserved even when individually silencing the i1, i2, b3, b1, or b2 motor neurons. Previous results showed that activation of b1 and b2 increases the stroke amplitude (***Whitehead et al., 2022***). We performed additional free flight optogenetic experiments to show that i1, i2, and b3 can decrease stroke amplitude and that i1 and i2 can decrease the wing pitch angle. Next, using quasi-steady aerodynamics we showed that stroke amplitude changes primarily affect z torque, while wing pitch changes primarily affect roll torque. Moreover, the corresponding changes during optogenetic manipulations can also produce torques consistent with those seen during roll corrections, and multiple muscles in each grouping can produce such torques. Analysis of the BANC connectome confirmed prior results found in the FANC connectome that these motor neurons separate into two groups based on inputs from the halteres: i1, i2, and b3 comprise one group, while b1 and b2 comprise the other (***Dhawan et al., 2025***).

We combined these results into a proposed schematic for how shared haltere inputs and redundant muscle outputs lead to robust roll and z torque production during roll maneuvers (***Figure 6)***. First, haltere sensory inputs separate into two clusters targeting different sets of muscles. Within each cluster, the constituent muscles are able to produce similar kinematic outputs: increasing stroke amplitude, in the case of b1 and b2, and decreasing stroke amplitude and wing pitch, in the case of b3, i1, and i2. These kinematic outputs then combine with each other to produce the desired corrective torques. In this scheme, even if one muscle is inhibited, there is always at least one other muscle producing the same kinematic output that can compensate for it, so the final roll and z torques are, to first order, unaffected.

**Figure 6.**
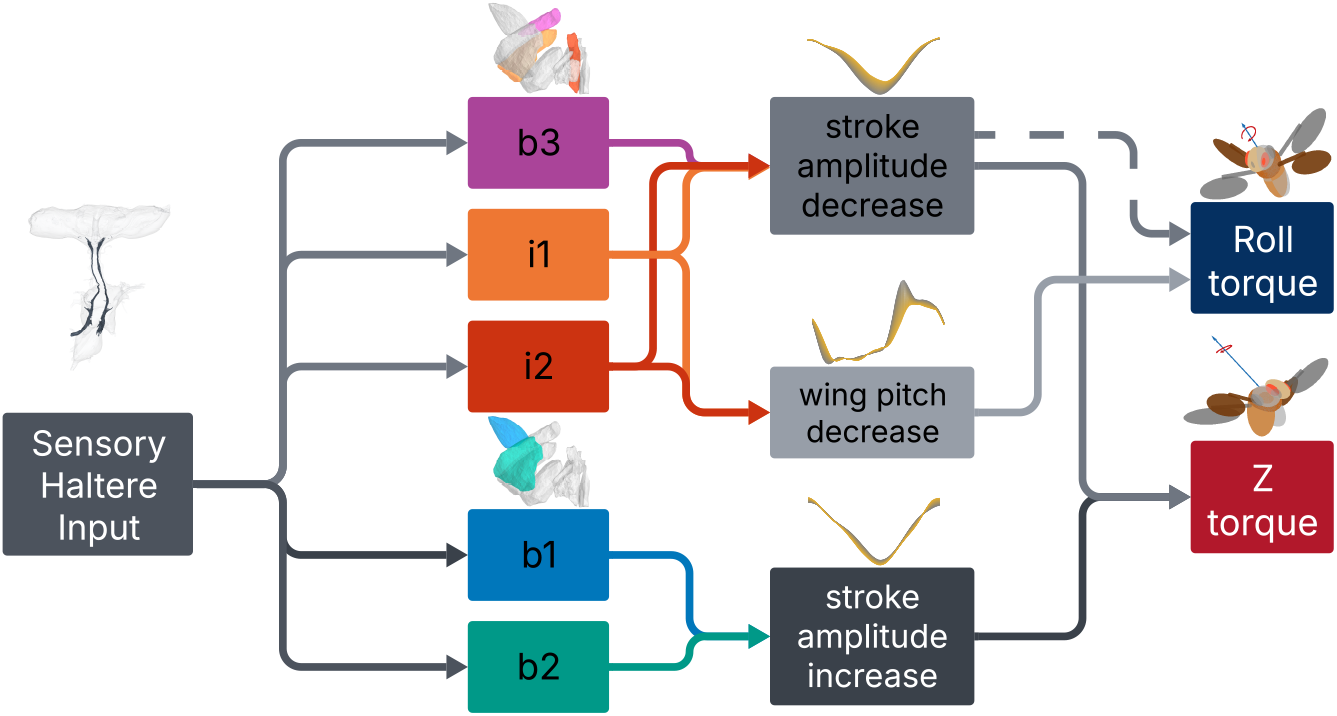
Proposed schematic of redundancies within the roll control system. Sensory haltere input is divided between two clusters of muscles based on their kinematic output. These kinematic changes then combine to produce corrective roll and z torques.

The robustness and possible redundancy within the neuromuscular implementation of roll control is notably different from previous results studying pitch, the fly’s other unstable rotational degree of freedom. In ***Whitehead et al. (2022)***, it was found that two of the basalar motor neurons implemented a significant portion of pitch control. Silencing the tonic b1 muscle removed the integral term of the controller for pitch, and silencing the phasic b2 muscle greatly diminished the proportional term in the pitch controller. Several analogous traits between the basaslar muscles and the other three muscles investigated in this paper—i1, i2,and b3—made them appear to be good candidates for implementing roll control in a similar way: i2 and b3 are both tonic muscles, while i1 is a phasic muscle; i1 and i2 both pull synergistically with each other, and along with b3 were hypothesized to be able to decrease stroke amplitude in similar ways (***Ehrhardt et al., 2025; Melis et al., 2024***); all three muscles showed qualitatively similar changes in their activity during visual roll perturbations in tethered flight (***Lindsay et al., 2017***). Therefore it was somewhat surprising that individually inhibiting the i1, i2 or b3 motor neurons did not significantly impact the fly’s ability to control for roll. Previous work had already shown that inhibition of b1 or b2 motor neurons did not affect the roll controller (***Whitehead et al., 2022***). Here, we expanded on these results to show that individual inhibition of these motor neurons also does not affect the relevant wing kinematics during roll corrections.

Prior work in other model systems has shown compellingly how preservation of function during the ablation of one or more sensorimotor systems does not necessarily imply that that apparatus is not used for the chosen task. For example, ***Roth et al. (2016)*** identified parallel visual and mechanical pathways used in hawkmoth flower tracking. In their study, hawkmoths were able to robustly track flowers in the dark, or alternatively in the absence of mechanosensory input, by adaptively tuning the gain of each sensory pathway to different sensory contexts. It was only by placing the two sensory modalities in conflict with each other that they were able to separate out the contributions of the two redundant systems. Similarly, in our system, although inhibiting individual motor neurons during roll perturbations had no impact on their corrections, such muscles could still be active during roll corrections in a redundant control architecture.

Redundancy is not without precedence in the Drosophila flight control architecture, too. In the previous work on the role of the basalar muscles in pitch control, while they were less stable, flies were still able to fly when the b1 or b2 muscles were silenced—a significant finding especially in the case of b1, which was long predicted to be essential for flight control (***Chang and Wang, 2014***). Additionally, while ablating the function of the b2 muscle reduced the proportional term in the pitch controller, it did not fully eliminate it (***Whitehead et al., 2022***). Further studies of pitch control have also found that flies make use of an additional kinematic strategy during large perturbations, altering their angle of attack to generate additional drag-based torques and minimize power consumption during corrective maneuvers. These changes are largely governed by the indirect tergopleural (tp) steering muscles, which had not previously been implicated in flight control (***Cohen et al., 2024***). Such findings could be evidence for a more complex architecture than has previously been laid out in pitch control, as well.

We hypothesize that the origin of necessity for redundancy in the neuromuscular control of roll lies in the degree of instability the fly experiences about the body roll axis, as well as the importance of roll for in-flight maneuverability. Of all the rotational degrees of freedom of the fly, roll has the shortest time scale of instability. Computational fluid dynamics calculations and aerodynamic simulations of stability dynamics in fruit fly flight have found that roll control requires a minimum response time of only 2 wingbeats, consistent with experimentally measured control dynamics (***Perl et al., 2023; Chang and Wang, 2014; Ristroph et al., 2013***). This constraint puts significant pressure on the muscular implementation of roll control to be fast, robust, and flexible. Tonic muscles such as b1, i2, and b3 have been hypothesized as the main pathway for this wingbeat-to-wingbeat control (***Chang and Wang, 2014***). However, because it has been shown that flies can still fly and maintain roll stability with any of these individual tonic muscles silenced, we propose a system in which multiple muscles are controlled together redundantly, to maintain stability even in the face of loss of function of any single muscle.

While the focus on flight stability seems to narrow our scope to only one facet of flight behavior, the control loop hypothesis posits that these same stability reflexes may be co-opted to drive volitional maneuvers via active manipulation of the haltere (***Verbe et al., 2024; Dickerson et al., 2019***). Thus, this redundancy may be important not only for stability, but also evasive and navigational saccades, of which roll is already to known to be a principle part of the maneuver (***Muijres et al., 2015***). As more and more powerful genetic and connectomic tools become available to Drosophila research, we hope the results presented here of how distinct muscles can combine redundantly for robust roll control will contribute to eventually decoding the neural architecture of insect flight behavior more broadly.

## Methods and Materials

### Fly stocks and fly handling

Flies used for optogenetic experiments were reared in the dark at room temperature on 0.4 mM retinal food (Media Facility, HHMI Janelia Research Campus) for a minimum of three days after eclosion. Flies used for all other experiments (e.g., mechanical perturbation) were raised at room temperature on standard fly medium made from yeast, agar, and sucrose with a 12-hour light/12- hour dark cycle. Female flies, 3 to 6 days after eclosion, were used for all flight experiments. A full list of Drosophila melanogaster stocks used in this paper is given in ***Table 1.***

**Table 1:**
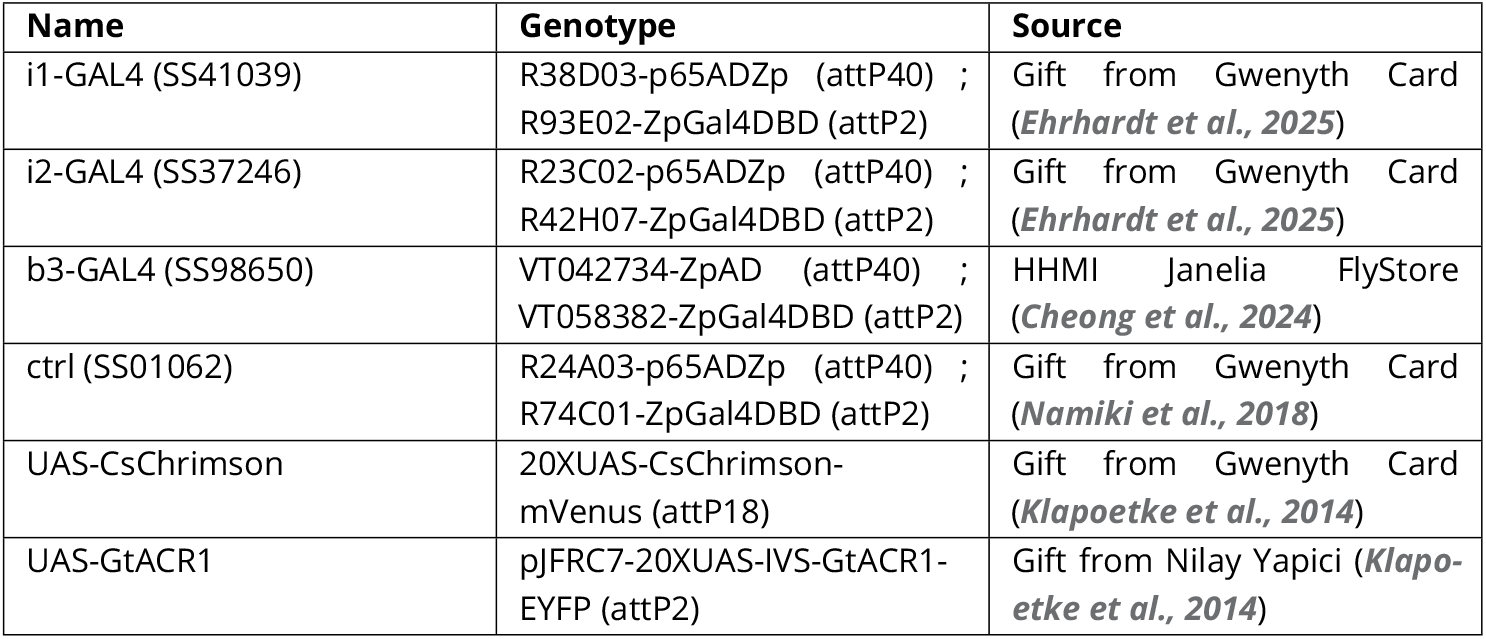
Fly stocks.

### Fly preparation

For mechanical perturbation experiments, we anesthetized individual flies at 0^°^ to 4^°^. We then glued 1.1- to 2.0-mm-long, 0.15-mm-diameter ferromagnetic pins their notum (dorsal thoracic surface). The pins where oriented perpendicular to the flies’ sagittal plane. Experiments using unpinned flies showed no qualitative change in the wing kinematics with the addition of the pins (***Ristroph et al., 2010***). The mass of the pins is comparable to intra-fly variation on body mass, and the pins add negligibly to the off-diagonal components of the fly’s inertia tensor (for detailed calculations, see ***Whitehead et al. (2015); Beatus et al. (2015)***).

### High-speed videography

We performed experiments with 10 to 30 flies prepared as above, all with uniform genotype. Flies were released into a transparent cubic flight chamber with a side length of 13 cm. The center of the chamber was filmed with three high-speed cameras (Phantom V7.1 or Phantom VEO 710) at 8000 frames/s and 512×512 pixel resolution, with a mutual filming volume of approximately 8cm^3^ (see ***Figure 1A***). Each camera was backlit by a focused 850- ± 30-nm near-infrared light-emitting diode (LED) (Osram Platinum Dragon). An optical trigger—created using split, expanded beams from a 5-mW, 633-nm HeNe laser (Thorlabs, HRR050) passed through a neutral density filter (Thorlabs, NE20A) with an optical density of 2.0 and incident upon two photodiodes (Thorlabs, FDS100)—was used to detect the entrance of flies into the filming volume of the high-speed cameras during experiments and initiate filming (***Ristroph et al., 2009***). Before each experiment, the cameras were calibrated using the easyWand system in (***Theriault et al., 2014***).

### Mechanical perturbation experiments

For each mechanical perturbation experiment, 10 to 25 flies were prepared by gluing small ferromagnetic pins to the dorsal side of their thoraces (see above) and subsequently released into the flight chamber. Similar to the optogenetic experiments described above, an optical trigger circuit was used to apply an 8-ms magnetic field pulse whenever a flying fly entered the center of the filming volume. High-speed cameras were simultaneously triggered to record flight activity before, during, and after the application of the magnetic field at 8000 frames/s, as described above.

The impulsive magnetic field was generated by the optical triggering circuit driving a rapid current pulse to two Helmholz coils mounted on the top and bottom faces of the flight chamber. Because of the positioning of the Helmholz coils, this generated an approximately uniform vertical magnetic field within the filming volume. Typical magnetic field strengths were on the order of approximately 10^−2^ T. The magnetic field acted on the magnetic moment of the ferromagnetic pin glued to the fly, generating a torque about the fly’s roll axis as the pin rotated to align with the magnetic field. Further details of this procedure are described in ***Ristroph et al. (2010); Whitehead et al. (2015); Beatus et al. (2015)***.

We performed these experiments in conjunction with optogenetic silencing. To do so, experiments were performed as described for the free-flight optogenetic experiments (see below), with the addition of ferromagnetic pins glued to the flies’ thoraces (see above) and a magnetic impulse. The timing of the optogenetic stimulus and the magnetic impulse were offset such that the optogenetic stimulus was delivered from t = 0 to 50 ms, and current was run through the Hemlholz coils from t = 15 to 22 ms. This resulted in a period of 15 ms of optogenetic silencing, a 7ms magnetic impulse under continued silencing, and then 28 ms of continued optogenetic silencing while the fly recovered from the mechanical perturbation. A version of this procedure has previously been used for pitch perturbations in ***Whitehead et al. (2022)***; **?**.

Wing kinematic changes during roll corrective maneuvers, as in ***Figure 1F***-H,J-L,O-R, were defined comparing “pre-perturbation” wingbeat to a “peak corrective” wingbeat. The pre-perturbation wingbeat was defined as the last wingbeat before (but not including) optogenetic stimulus onset. The peak corrective wingbeat was defined as the wingbeat during the correction period (t = 22 to 50 ms) with the minimum stroke amplitude on the upper wing. This heuristic was chosen as the upper stroke amplitude decrease was the most robust of the measured correction mechanisms, and the chosen wingbeats tended to show good agreement with peak corrective roll accelerations.

### Flight data selection and kinematic extraction

When analyzing the data for the free-flight optogenetic, mechanical perturbation, and combined optogenetic and mechanical perturbation experiments described above, we restricted our analysis to flight bouts which were amenable to kinematic extraction. For free-flight optogenetic experiments, this criterion meant the fly would need to be in full view of all three cameras for a period of at least 10 ms prior to the onset of the optogenetic stimulus, and at least 45 ms after stimulus onset. Additionally, we excluded videos in which the fly was not flying stably at the time of stimulus onset (eg, fly was falling rather than flying, or was experiencing uncontrolled midair rotations prior to any stimulus). For experiments which included a mechanical perturbation (as in ***Figure 1,*** we used additional stricter criteria. We only used videos in which the perturbation was principally about the body roll axis, rather than a mix of roll and pitch or roll and yaw. For ***Figure 1T***-V this also meant limiting our data to cases in which fits to our proportional-integral controller gave physically reasonable results (see Controller model fitting for single-trial data, below). These restrictions were necessary to ensure the kinematics extracted after the mechanical perturbation could be cleanly attributed to corrective maneuvers about a single body axis of rotation.

To extract wing and body kinematics from our three high-speed videos, we used a custom hull reconstruction algorithm detailed in ***Ristroph et al. (2009)***. Using this algorithm, we obtained a 12 degree-of-freedom description of the fly—the 3D position of the fly center of mass and the three full sets of Euler angles for the fly body, left wing, and right wing—at each time point. For time points in which occlusion prevented the direct extraction of a particular kinematic variable, we used a cubic spline interpolant to fill in missing data values. For most analyses, raw body kinematics were filtered using a 100-Hz low-pass filter. Raw wing kinematics were smoothed using the Savitzky-Golay method. For the wing stroke angle, we used a polynomial order seven with a window size of 21 frames (2.625 ms); for the wing deviation and rotation angles, we used a polynomial order five and a window size of 11 frames (1.375 ms).

To average wingbeat kinematics across flies—as in ***Figure 1F***-L,O-R; ***Figure 2B***,C,E,F,H,I; ***Figure 3B***; and ***Figure 4B***—we segmented wingbeats from time series data based on back wing flip times, i.e., the frame in which the wing pitch angle crossed 90^°^ while the wings were at the rearmost stroke position. Segmented wingbeat kinematics were then aligned to nondimensional wingbeat cycle time using a sixth-order Fourier series fit (MATLAB’s fit.m using the “fourier6” option) to evenly resample the Euler angle values. With all segmented wingbeats sampled according to a common nondimensional time, wing Euler angle traces could be directly averaged without incurring errors because of varying wingbeat frequency across flies/wingbeats.

### Immunohistochemistry

To visualize the nervous system and expression patterns of our split-GAL4 driver lines (as in ***Figure 1N***), we followed a standard VNC imaging protocol based on those used in ***Whitehead et al. (2022)***; **?**. We dissected *drosophila* CNS in phosphate-buffered saline (PBS) using fine forceps and placed them in 4% paraformaldehyde solution (in PBS) for 15 min at room temperature. Then, we washed the CNS in phosphate-buffered saline with Tween®(PBST) three times, 15 minutes each, before incubating the CNS in normal goat serum (NGS) for 20 minutes. We stained the tissue overnight at 4^°^ C with primary antibodies consisting of 1:10 mouse anti-nc82 (DSHB) and 1:1000 rabbit anti-GFP (Invitrogen no. A6455) in PBST. Next, we washed the CNS in PBST and applied a secondary antibody stain consisting of 1:250 goat anti-mouse AlexaFluor 633 (Thermo Fisher no. A21052) and 1:250 goat anti-rabbit AlexaFluor 488 (Thermo Fisher no. A11034) in PBST overnight at 4^°^ C. We then pipetted the CNS from the solution and placed them in PBS for 30 min on an oscillator at room temperature. Finally, we mounted the CNS on a glass microscope slide in Vectashield (Vector Labs H-1000-10), covered with one #1.5, 22 × 22 mm coverslip, and sealed with nail polish. We imaged the CNS at 20x magnification on a Zeiss LSM 880 confocal microscope at 1 *µ*M depth resolution. Final images are presented at maximum z-projections over relevant depths.

### Controller model fitting for single-trial data

For each movie of a fly performing a corrective maneuver after a mechanical perturbation (see ***Figure 1S***-V), we fit a proportional-integral (PI) controller model to the kinematic data obtained as described above. For roll corrective maneuvers, the PI controller is governed by the following equation:

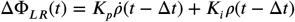

where ΔΦ_*LR*_ is the difference between left and right stroke amplitude, *ρ* is the body roll angle, and *K*_*p*_ and *K*_*i*_ are the proportional and integral controller gains, respectively. This model is based on the one previously derived in ***Beatus et al. (2015)***. Note that the sign convention for this equation is chosen such that *K*_*p*_ and *K*_*i*_ are positive for a stable system, as in ***Beatus et al. (2015)***.

To fit the controller gains from our experiments, we extracted Φ_*LR*_ in the time period of the corrective maneuver from each of our movies. We performed a nonlinear least squares fit (Levenberg-Marquardt) for the gain coefficients, *K*_*i*_ and *K*_*p*_, along with a grid search for the time delay, Δ*T* . In some cases, the grid search failed to find a minimum for Δ*T* within physically reasonable bounds; we set a minimum value of 2 ms based on previous estimates of anatomical limits on the fruit fly’s reaction time (***Chang and Wang, 2014; Ristroph et al., 2013)***. These cases were excluded from our analysis as they did not provide an accurate measurement of the controller parameters. Using this procedure, we obtained three parameters from each movie which could model the fly’s full corrective response: *K*_*p*_, *K*_*i*_, and the time delay Δ*T* . Uncertainty in fit parameters was estimated using the fit parameter covariance matrix at the objective function minimum. The covariance matrix *C* was approximated as *C* ≈ *σ*^2^(*J* ^*T*^ *J*)^−1^, where *σ*^2^ is the variance of the fit residuals and *J* is the calculated Jacobian at the objective function minimum. The within-genotype spread of controller parameters seen in ***Figure 1T***-V is attributed to a combination of variations in morphology—across individuals and also overtime as flies were more or less hydrated during the course of an experiment—as well as noise injected into the system by fitting time-domain models to perturbation events which occur over relatively short time windows.

### Free-flight optogenetic experiments

For free-flight optogenetic experiments, 10 to 30 flies were released into the flight chamber described above for a period of approximately 12 hours. To apply midflight optogenetic manipulation, we used the optical trigger system described above to deliver a 50-ms light pulse from a collimated LED source located outside the chamber (***Figure 1A***) whenever the fly entered the center of the filming volume. This trigger simultaneously initiated filming by the three high-speed cameras, which captured flight behavior before, during, and after the light stimulus as described above. To apply the light stimulus, the optical trigger circuit drove a 50-ms duration voltage pulse to an LED driver (Thorlabs, LEDD1B), which was connected to either a 625-nm red LED (Thorlabs, M625L4) or a 565-nm green LED (Thorlabs, M565L3) for optogenetic activation (CsChrimson) or inhibition (GtACR1) experiments, respectively. Both red and green LED sources were outfitted with a collimating attachment (Thorlabs, COP2-A) to generate a 50-mm-diameter beam profile. The beam diameter was large enough such that a fly located anywhere in the filming volume would experience the optogenetic stimulus, and the collimating attachment ensured uniform stimulus intensity throughout the entire filming volume.

The stimulus LEDs were driven with a 1-A current, which resulted in intensities of 731 and 316 *µ*W/mm^2^ for the red and green LEDs, respectively. Despite the optical trigger’s 633-nm HeNe laser ostensibly falling in the wavelength range of CsChrimson sensitivity, the optical filters on this light source ensured that the laser’s intensity, approximately 0.16 *µ*W/mm^2^, was two to three orders of magnitude lower than any applied LED stimulus. Moreover, experiments performed in ***Whitehead et al. (2022)*** found no changes in flight behavior due to the triggering laser when LED stimulus was withheld.

To prevent contamination from outside light sources, the entire apparatus was enclosed in blackout curtains during experiments. Because flies are less likely to initiate flight bouts in complete darkness, a dim, blue fluorescent light bulb was used to illuminate the chamber during experiments.

To analyze the flight kinematics from these experiments, “pre-stim” and “stim” periods—as in ***Figure 2—***were defined relative to the onset of the LED stimulus, to capture data before and during optogenetic manipulation. For analysis performed in ***Figure 2,*** the four wingbeats prior to the onset of the LED stimulus, but not the wingbeat during onset, were defined as the pre-stim period. The fourth to seventh wingbeats after LED onset, while the LED was still on, were defined as the stim period. The results of the optogenetic stimulus were not sensitive to the particular wingbeat selected, and the selection of wingbeats for stimulus period was based on previous analysis in ***Whitehead et al. (2022)***.

### Quasi-steady analysis procedure and flight simulation parameters

For the analysis performed in ***Figure 3***and ***Figure 4,*** we selected a set of representative movies for each condition (magnetic perturbation of optogenetic controls for ***Figure 3,*** and i1, i2, b1, b2 stimulation and b3 inhibition for ***Figure 4)*** from the data presented in ***Figure 1E***-L and ***Figure 2.*** Our selection criteria were (1) the fly was in full view of all three cameras from t = -20 ms to t = 50 ms, (2) the fly did not perform any volitional maneuvers during that same time period, and (3) the wing kinematics observed in the movie provided a qualitatively representative example of the average wing kinematics across the population of experiments. This was done to ensure that there was sufficient data to capture the time course of the kinematic changes during roll perturbations and/or optogenetic manipulation, and that the wing kinematic changes—and resulting torques— could be attributed to the effects of the perturbations.

For each movie, we selected one “pre-perturbation” wingbeat (defined as the last full wingbeat before the onset of the optogenetic stimulus) and one “maximally corrective” (***Figure 3)*** or “post-stimulus” (***Figure 4)*** wingbeat. We segmented wingbeats from time series data based on back wing flip times, i.e., the frame in which the wing pitch angle crossed 90^°^ while the wings were at the rearmost stroke position. The “maximally corrective” wingbeat was defined as the wingbeat with the greatest decrease in upper stroke amplitude. The “post stimulus” wingbeat was a wingbeat after 30-40 ms of optogenetic stimulation, chosen by hand to ensure clean wing kinematic reconstruction as well as a clear phenotype from the optogenetic stimulus. To generate the 50 wing kinematic profiles for each kinematic variable shown in ***Figure 3B***, ***Figure 4B***, ***Figure 4—figure Supplement 1***, ***Figure 4—figure Supplement 2***,***Figure 4—figure Supplement 3***, and ***Figure 4—figure Supplement 4***, we used a 1-D data interpolation algorithm (MATLAB’s interp1.m function) to align each wingbeat to a common non-dimensional time vector with *N* = 50 points while smoothly sampling the kinematic variables over that interval. This allowed us to generate linear combinations of the wingstroke profiles from before and during stimulation without causing misalignment errors due to variation in wingbeat frequency over the course of the trial. We then evenly sampled a set of 50 linear combinations varying from 100% pre-perturbation kinematics to 100% maximally corrective/post-stimulus kinematics for each kinematic variable.

We selected combinations of kinematic profiles which allowed us to independently change one kinematic variable while co-varying the other five (***Figure 3)*** or two (***Figure 4)*** variables. For each set of profiles, the net roll and z torques were calculated at each time point across our sample using the quasi-steady aerodynamic simulations reported on in ***Whitehead et al. (2015, 2022); Dickinson et al. (1999); Sane and Dickinson (2002)***. The total instantaneous quasi-steady aerodynamic force on the wing, **F**, is separated into translational (**F**_**t**_) and rotational (**F**_rot_) terms (***Dickinson et al., 1999; Sane and Dickinson, 2002; Whitney and Wood, 2010; Dickinson and Muijres, 2016)***; we exclude the contribution of added mass in this model, following ***Whitehead et al. (2022)*** and ***Whitney and Wood (2010)***, as analytic expressions for its contribution are inaccurate in cases of large acceleration. Thus, the total force is,

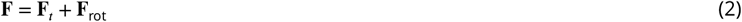

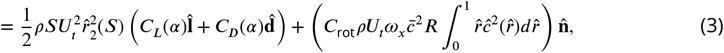

Where *ρ* is the air density; *S* is the wing area, approximated as the area of an ellipse; *U*_*t*_ is the wing tip velocity; 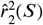 is the non=dimensionalized second moment of wing area; *α* is the wing angle of attack; *C*_*L*_ and *C*_*D*_ are the lift and drag coefficients, respectively; *C*_rot_ is the coefficient of rotational force; *ω*_*x*_ is the angular velocity of the wing about its spanwise axis; 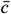 is the mean wing chord length; *R* is the wing span length; 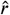 and 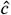 are the non-dimensionalized span and chord; and 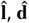, and 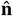 are unit vectors in the direction of the lift, drag, and wing surface normal, respectively. The lift and drag coefficients are further expressed as functions of the wing angle of attack (***Wang et al., 2004)***:

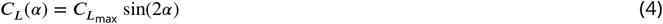

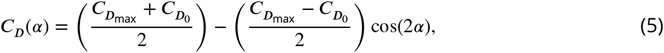

Where 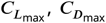, and 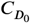 are dimensionless constants that have been best fit for *Drosophila* flight.

Finally, the aerodynamic torque **T** is calculated by approximating that the wing center of pressure is located 70% of the way along the length of the wing span (***Birch and Dickinson, 2001)***:

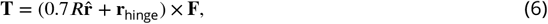

where *R* is the wing span length; 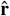 is the wing span unit vector; **r**_hinge_ is the vector from the fly’s center of mass to the wing hinge; and **F** is the total aerodynamic force on the wing. We then averaged these torques over the entire wingbeat and repeated this procedure across many combinations of kinematic profiles to generate the heatmaps shown in ***Figure 3C***,E-G,I-K and ***Figure 4C***-Q. A full set of simulation constants and parameters is given in ***Table 2.*** Constant values for the morphological and physical parameters were used for all flies.

**Table 2:**
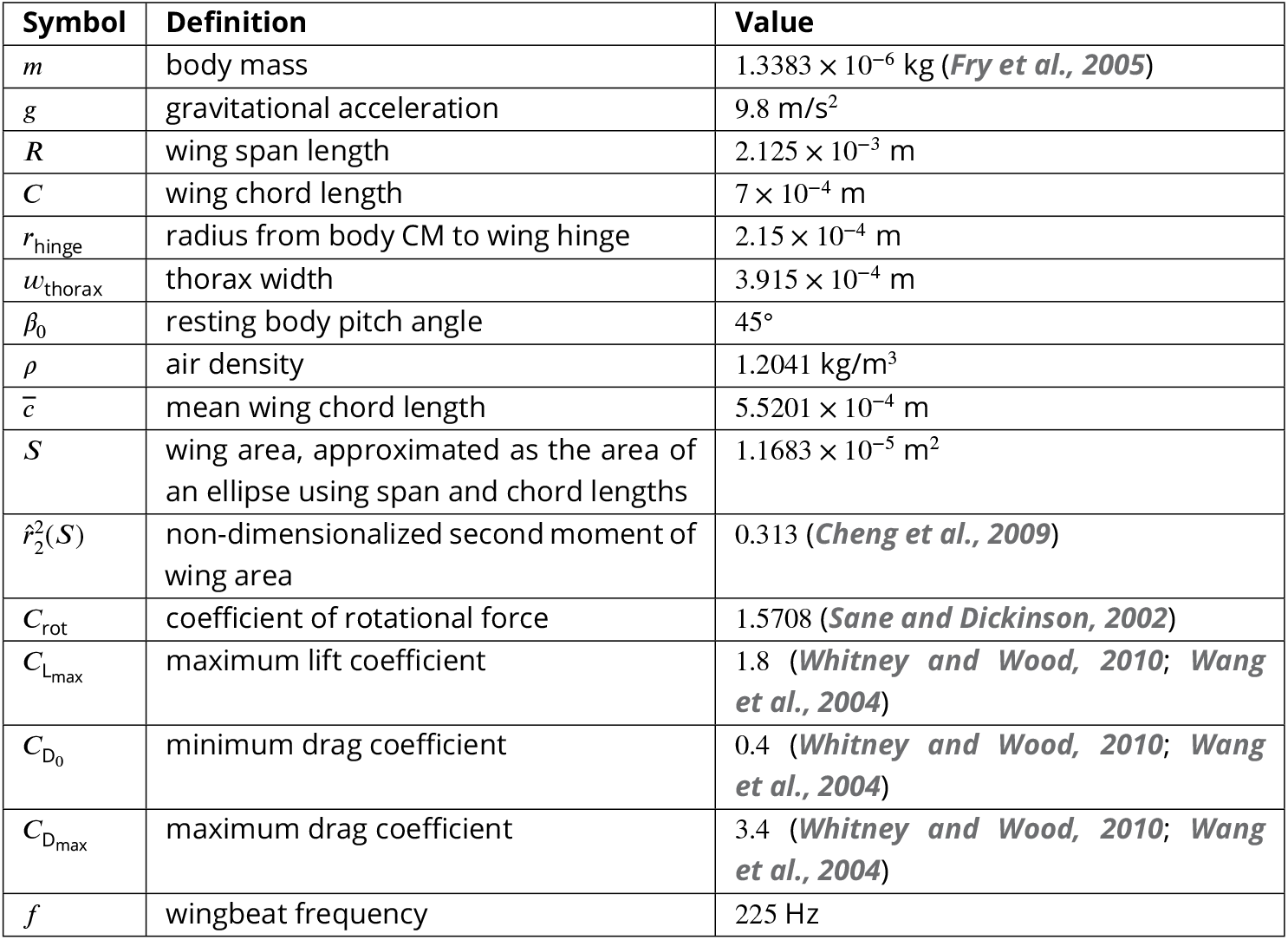
Simulation constants and parameters.

## Acknowledgments

We thank Gwenyth Card for providing the SS41039, SS37246, SS01062, and UAS-CsChrimson lines, and we also thank Nilay Yapici for providing the UAS-GtACR1 line. We are gratefult to Marie Suver’s lab for providing the protocol for CNS dissection and feedback and guidance along the way with that process.

---

**Figure 1—figure supplement 1.**
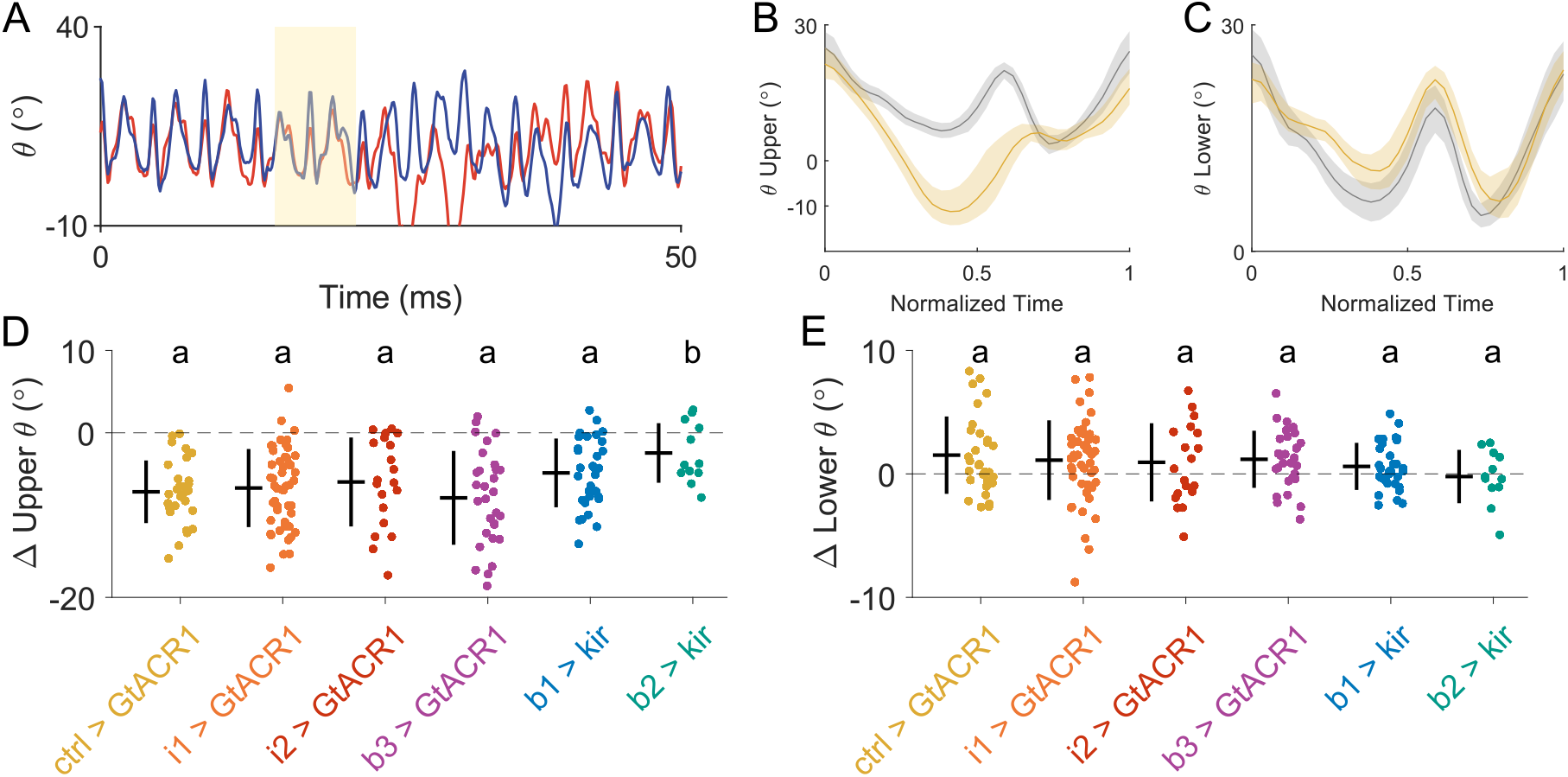
Deviation angle changes during roll perturbations. **A** Deviation angle of the left (blue) and right (red) wings over time during a roll perturbation. **B** Deviation angle of the uppermost wing for a single wingbeat before (grey) and during (yellow) a roll correction, averaged across all movies of genetic control flies (SS01062 > UAS-GtACR1; N = 17). Shaded regions represent bootstrapped confidence interval. **C** As in B, for the lower wing. **D** Change in wingbeat-averaged upper wing deviation angle during peak correction across all movies with individual motor neurons inhibited (N = 30, 50, 20, 30, 31, and 12, respectively). Data for b1, b2 inhibition are from ***Whitehead et al. (2022)***. **E** As in D, showing changes in wingbeat-averaged lower wing deviation angle. All statistics are Kruskal-Wallis test with Bonferroni correction for multiple hypotheses.

**Figure 1—figure supplement 2.**
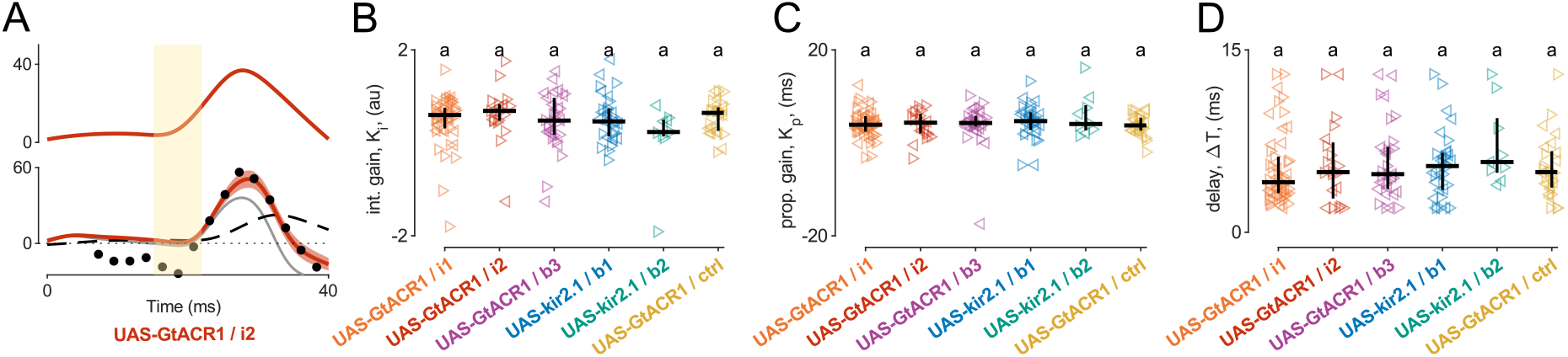
**A** Example roll controller fit. Top: Body roll angle over time during a roll perturbation. Yellow bar indicates magnetic impulse. Bottom: proportional (grey, solid) and integral (black, dashed) controller terms summed to predict stroke amplitude difference (red), compared to measured stroke amplitude data (black dots). **B** Measured integral gains across all movies with individual motor neurons inhibited (N = 50, 20, 30, 31, 9, and 30, respectively). Data for b1, b2 inhibition are from ***Whitehead et al. (2022)***. Statistics are Kruskal-Wallis test with Bonferroni correction for multiple hypotheses. **C** As in R, for measured proportional gains. **D** As in R, for measured time delays.

**Figure 2—figure supplement 1.**
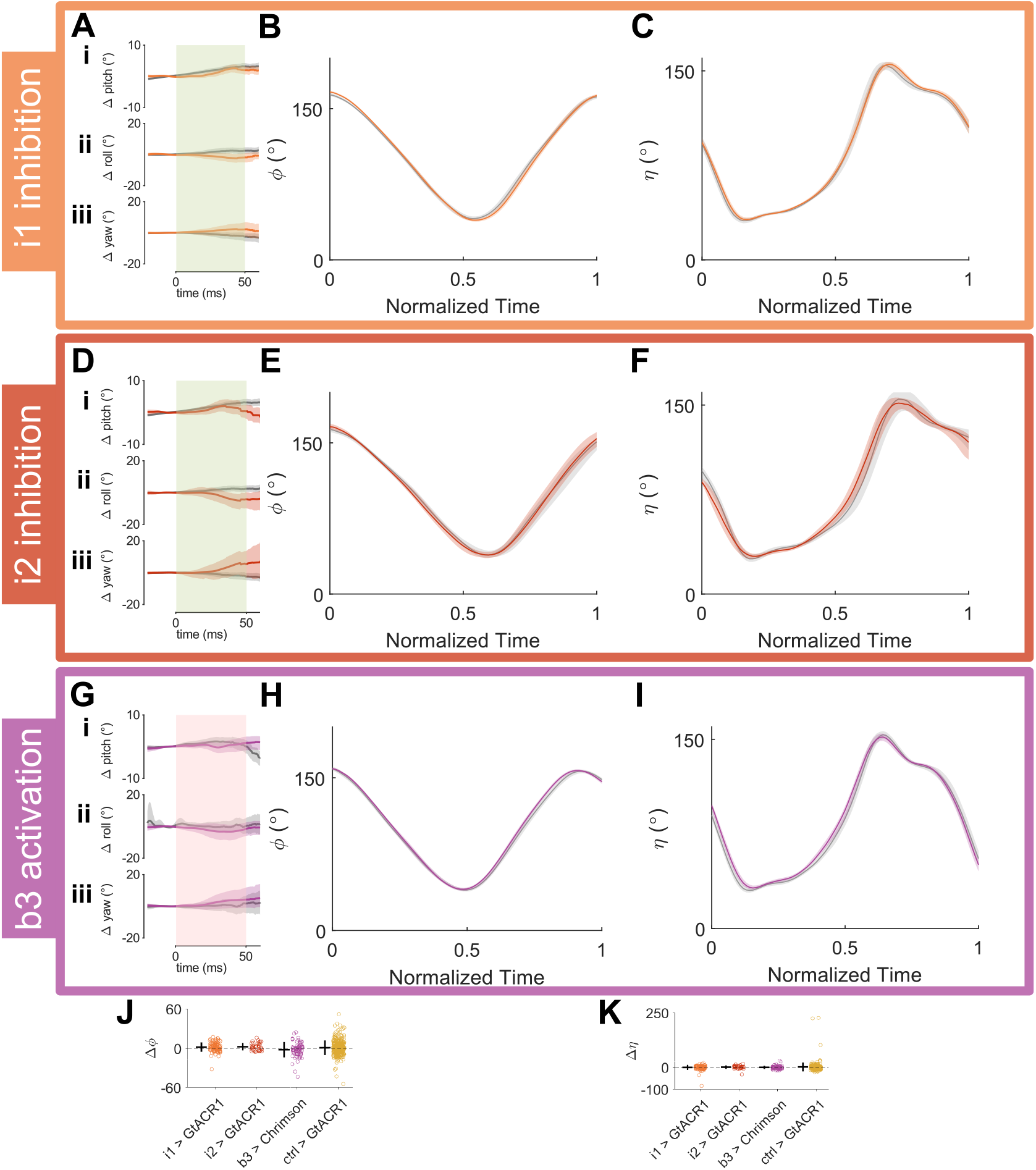
Optogenetic inhibition of i1, i2 and activation of b3. **A** Average body pitch (i), roll (ii), and yaw (iii) during optogenetic inhibition of the i1 motor neuron (SS41039 UAS-GtACR1, orange, N = 100) compared to genetic controls (SS01062 > UAS-GtACR1, grey, N = 345). Red bar indicates timing of optogenetic stimulus. Shaded areas represent bootstrapped confidence interval. **B** Stroke angle of for a single wingbeat before (grey) and during (orange) optogenetic inhibition of the i1 motor neuron, averaged across all movies. **C** Wing pitch angle for a single wingbeat before (grey) during (orange) optogenetic inhibition of the i1 motor neuron, averaged across all movies. **D-F** As in A-C, for optogenetic inhibition of the i2 motor neuron (SS37246 > UAS-GtACR1, red, N = 34). **G-I** As in A-C, for optogenetic activation of the b3 motor neuron (SS98650 UAS-CsChrimson, purple, N = 68. Genetic controls SS01062 > UAS-CsChrimson, grey (A i-iii), N = 195). **J** Change in stroke amplitude across all free-flight opotogenetic experiments. (N = 100, 34, 68, and 345 respectively). **K** As in J, for the change in peak wing pitch angle.

**Figure 3—figure supplement 1.**
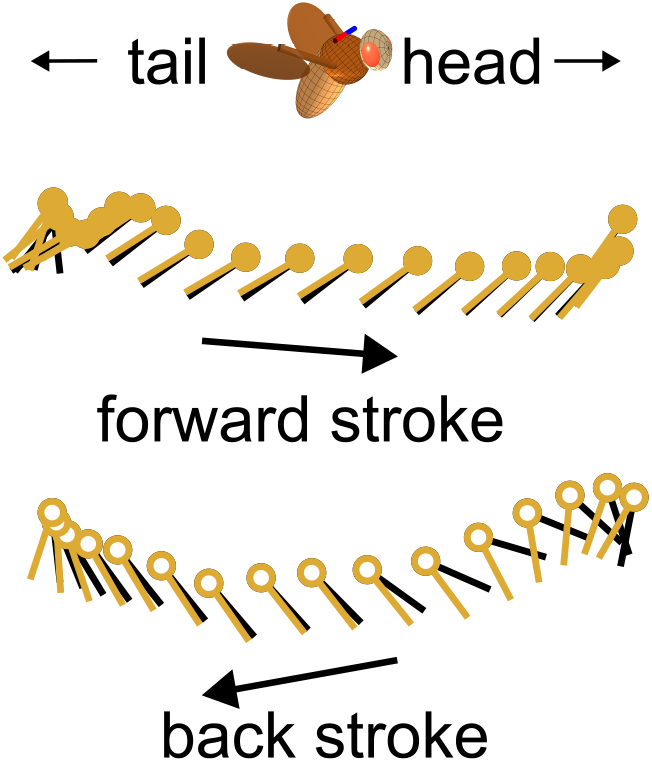
Visualization of changing only upper wing pitch during wingstroke. Top: Schematic illustrating direction of wingstroke. Middle: Ball and stick plot of the front stroke showing effect of wing pitch angle before and after a roll perturbation, with wing stroke and deviation angles keept at pre-perturbation trajectories. Bottom: As in the middle, for the back stroke. Wing trajectories are the same analyzed in ***Figure 3.***

**Figure 3—figure supplement 2.**
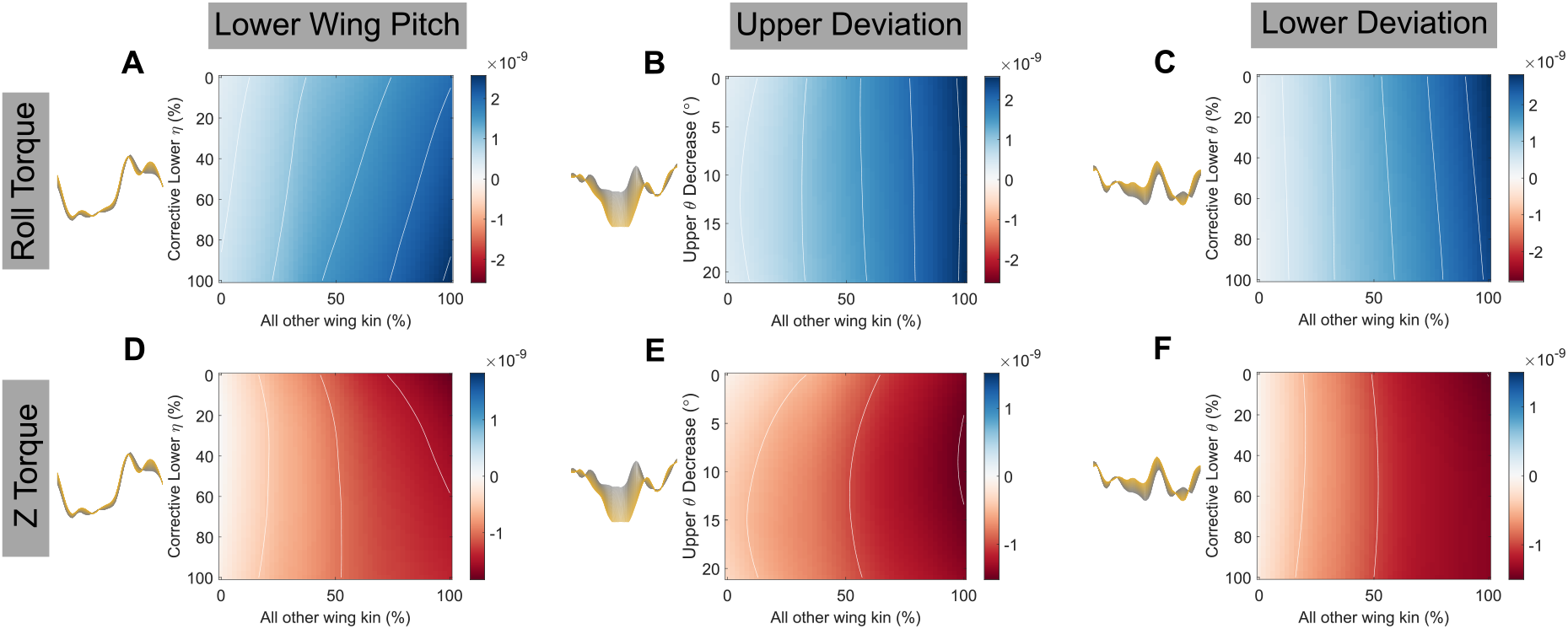
Quasi-steady aerodynamic calculations for other wing kinematic variables. **A** Wingbeat-averaged roll torque generated by different combinations of the wing stroke profiles in ***Figure 3***B. Y axis varies only lower wing pitch from pre-perturbation (grey) to maximally corrective (yellow). X axis uniformly varies all other wing kinematic profiles from pre-perturbation to maximally corrective. **B** As in A, with upper deviation profile varying along the y axis and all other kinematic profiles varying along the x axis. **C** As in A, with lower deviation profile varying along the y axis and all other kinematic profiles varying along the x axis. **D-F** As in A-C, with wingbeat average z torque plotted.

**Figure 4—figure supplement 1.**
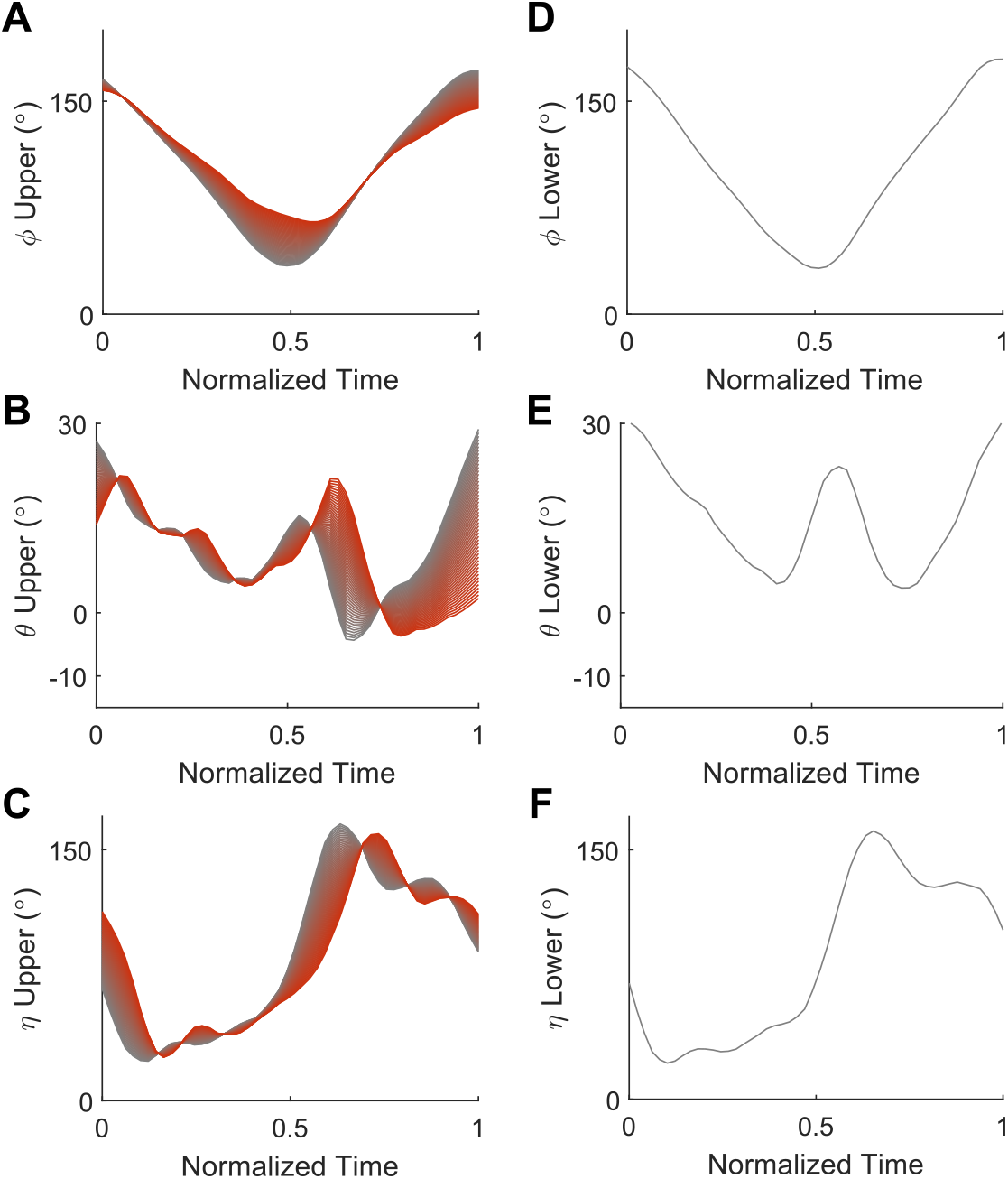
50 Linearly interpolated wing kinematic profiles for a single wing-beat, ranging from before stimulus (grey) to approximately 30 ms after i2 activation (red) on one wing, while the other wing kinematic profiles are kept static (pre-stimulus kinematics, grey): **A** “upper” (right) wing stroke angle, **B** “upper” (right) wing deviation angle, **C** “upper” (right) wing pitch angle, **D** “lower” (left) wing stroke angle, **E** “lower” (left) wing deviation angle, and **F** “lower” (left) wing pitch angle.

**Figure 4—figure supplement 2.**
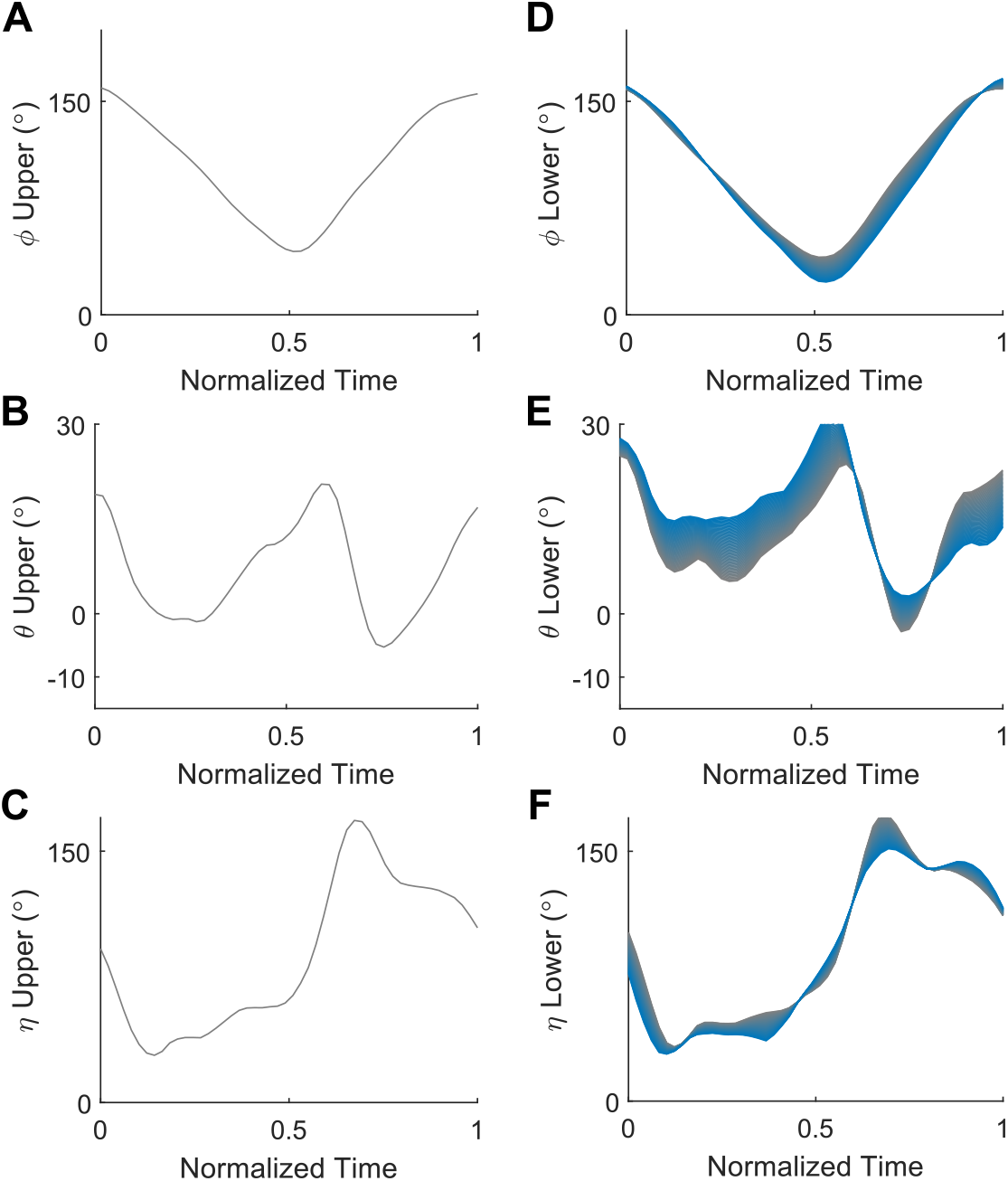
50 Linearly interpolated wing kinematic profiles for a single wingbeat, ranging from before stimulus (grey) to approximately 30 ms after b1 activation (blue) on one wing, while the other wing kinematic profiles are kept static (pre-stimulus kinematics, grey): **A** “upper” (right) wing stroke angle, **B** “upper” (right) wing deviation angle, **C** “upper” (right) wing pitch angle, **D** “lower” (left) wing stroke angle, **E** “lower” (left) wing deviation angle, and **F** “lower” (left) wing pitch angle.

**Figure 4—figure supplement 3.**
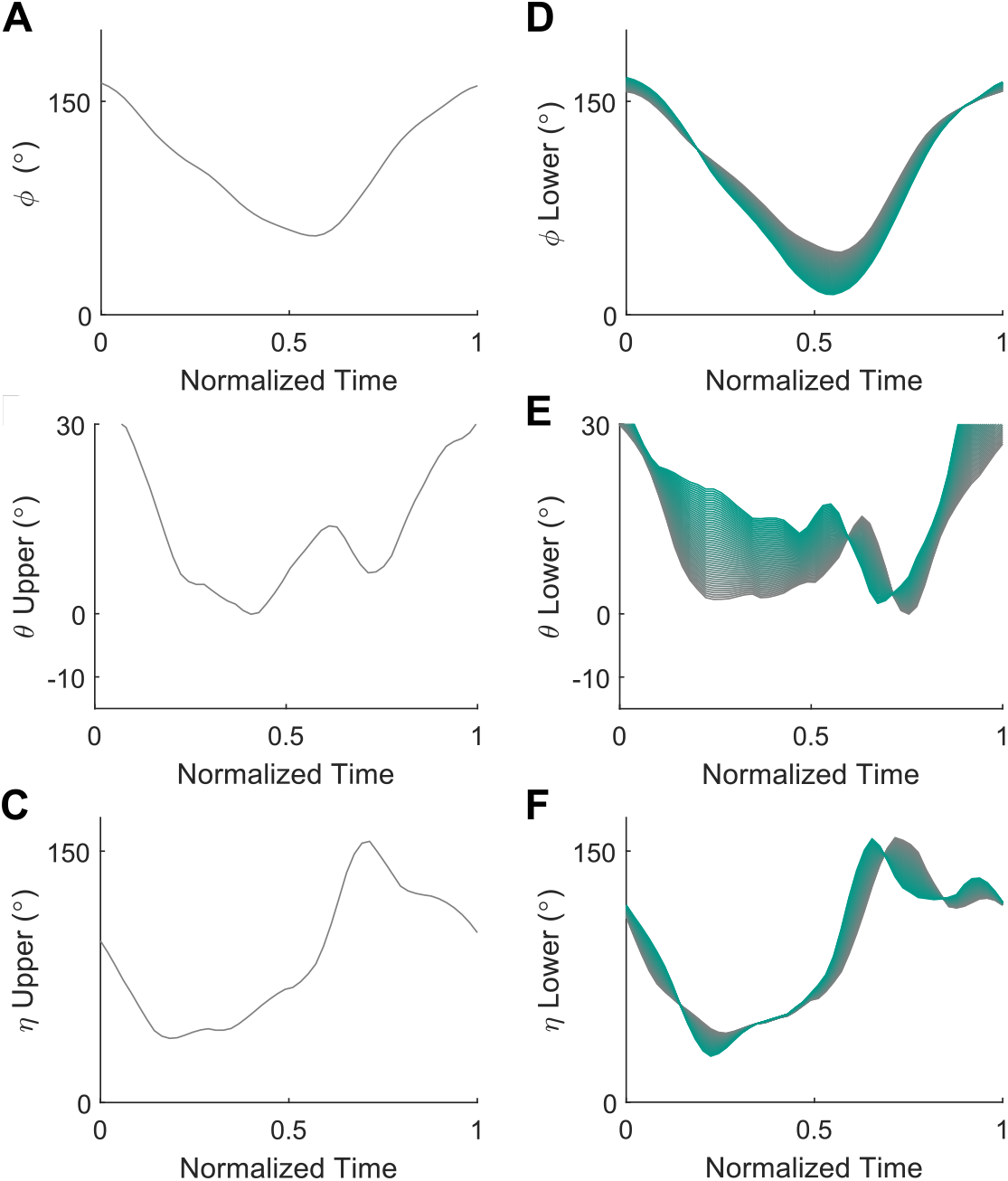
50 Linearly interpolated wing kinematic profiles for a single wing-beat, ranging from before stimulus (grey) to approximately 30 ms after b2 activation (teal) on one wing, while the other wing kinematic profiles are kept static (pre-stimulus kinematics, grey): **A** “upper” (right) wing stroke angle, **B** “upper” (right) wing deviation angle, **C** “upper” (right) wing pitch angle, **D** “lower” (left) wing stroke angle, **E** “lower” (left) wing deviation angle, and **F** “lower” (left) wing pitch angle.

**Figure 4—figure supplement 4.**
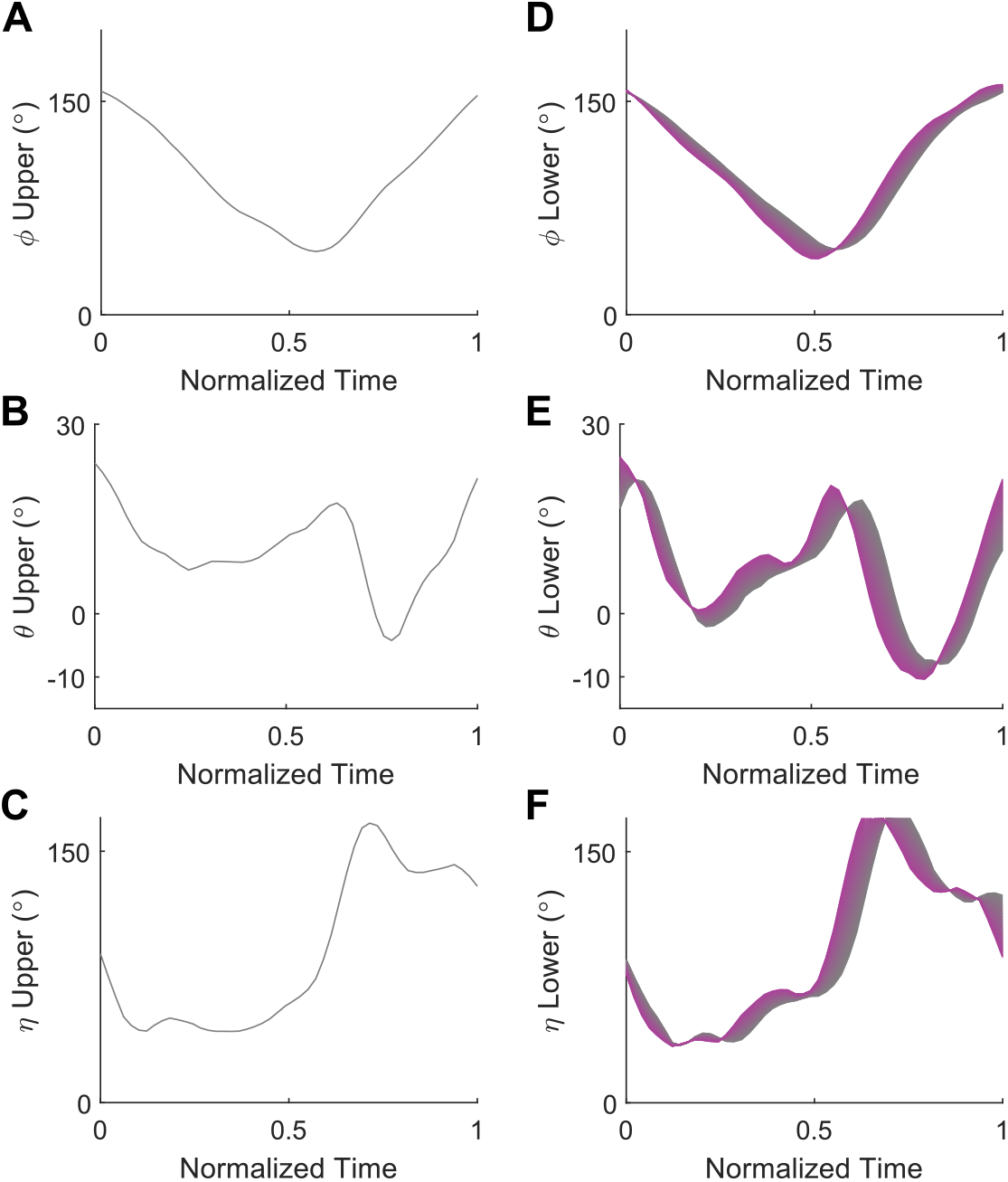
50 Linearly interpolated wing kinematic profiles for a single wing-beat, ranging from before stimulus (grey) to approximately 30 ms after b3 inhibition (purple) on one wing, while the other wing kinematic profiles are kept static (pre-stimulus kinematics, grey): **A** “upper” (right) wing stroke angle, **B** “upper” (right) wing deviation angle, **C** “upper” (right) wing pitch angle, **D** “lower” (left) wing stroke angle, **E** “lower” (left) wing deviation angle, and **F** “lower” (left) wing pitch angle.

